# Muscarinic signaling sustains prefrontal representations during short-term memory

**DOI:** 10.64898/2026.10.20.760355

**Authors:** Yeri Kim, Jinmo Kim, In-Sun Choi, Jungmin Lee, Yoon-Sil Yang, Joon Ho Choi, Ji-Woong Choi, Jong-Cheol Rah

## Abstract

Short-term memory (STM) requires the temporary maintenance of task-relevant information in prefrontal cortex (PFC). Reciprocal interactions between the mediodorsal thalamus (MD) and PFC are implicated in sustaining this activity, but how thalamofrontal communication persists despite constraints on repetitive synaptic transmission remains unclear. Here, we show that MD–PFC synapses exhibit profound frequency-dependent short-term depression, progressively reducing the ability of repetitive thalamic input to drive PFC neurons. During STM performance, acetylcholine (ACh) levels in PFC rapidly increased after cue presentation and remained elevated throughout the delay period. Muscarinic ACh receptor (mAChR) activation did not alleviate synaptic depression but enhanced intrinsic PFC neuronal excitability, preserving responsiveness to weakened thalamic input. Conversely, mAChR blockade impaired STM performance and progressively disrupted delay-period directional representations at both single-neuron and population levels, while largely sparing initial encoding. These findings identify muscarinic signaling as a postsynaptic gain mechanism that compensates for declining thalamofrontal efficacy and sustains task-relevant prefrontal representations during STM.

## INTRODUCTION

Short-term memory (STM) involves the temporary retention of task-relevant information, often attributed to sustained or dynamically evolving activity in the prefrontal cortex (PFC). While classical models emphasize persistent spiking as a substrate for memory maintenance ^1–3^, recent work in rodents and primates has highlighted alternative dynamics, including sequential firing, ramping activity, and low-dimensional population trajectories ^4–8^. Multiple long-range circuits have been implicated in shaping these PFC dynamics during STM. Among them, the thalamo-cortico-thalamic (TCT) loop ^6,9–11^, particularly the reciprocal excitatory connections between the mediodorsal thalamus (MD) and PFC, has emerged as a prominent candidate for stabilizing and enhancing memory-relevant representations ^10,12^. However, a recurrent excitatory loop alone does not explain how the efficacy of thalamofrontal transmission is dynamically regulated across the rapidly changing demands of memory encoding, maintenance, and termination.

Thalamocortical synapses are known for their high reliability and frequency-dependent short-term depression, which can act as a dynamic filter to regulate signal transmission depending on behavioral state ^13–15^ (but see ^16,17^). Consistent with the importance of maintaining thalamocortical transmission, we recently found that NMDAR hypofunction, a condition associated with STM deficits in schizophrenia, selectively reduced neurotransmitter release at thalamofrontal synapses, thereby further compromising thalamocortical information transfer ^18^. A key unresolved question in the TCT loop hypothesis is therefore how this circuit maintains the fidelity of information transfer in the face of synaptic depression and fluctuating cortical excitability.

STM depends critically on the sustained allocation of attention to task-relevant information ^19^ yet the precise mechanisms underlying this functional coupling remain poorly understood. Neuromodulators, including dopamine, norepinephrine, and acetylcholine (ACh), have long been implicated in shaping cortical function and cognitive performance. In particular, elevated PFC activity during STM is highly sensitive to changes in neuromodulatory tone in non-human primates ^20–22^. Among these modulators, ACh plays a prominent role in regulating attention, arousal, and learning and memory ^23–27^. In the PFC, ACh is primarily released by basal forebrain cholinergic projections ^28,29^ and enhances the intrinsic excitability of local excitatory neurons via muscarinic acetylcholine receptors (mAChRs) ^30^. Consistent with a broader role in cortical information processing, transient ACh release improves cue detection ^31^, and differentially regulates thalamocortical and intracortical synaptic transmission ^32–34^, suggesting that it may dynamically control cortical information flow during cognitive processing. Despite these findings, it remains unclear how ACh supports STM at the circuit level. In particular, whether cholinergic modulation preserves information flow through the TCT loop by compensating for the progressive loss of thalamofrontal efficacy during repetitive activity—and thereby sustaining thalamic drive during STM—remains an open question.

Here we tested the hypothesis that cholinergic signaling dynamically preserves information flow through the TCT loop by compensating for the loss of thalamofrontal efficacy during sustained activity. Combining *in vitro* electrophysiology, *in vivo* pharmacology, and neural population recordings in behaving mice, we show that muscarinic receptor activation maintains effective thalamofrontal transmission despite strong synaptic depression, thereby preserving delay-period representations and STM performance.

## Methods

### Animals and stereotaxic surgeries

For behavior, C57BL/6J (JAX: 000664) mice were used. The mice were housed individually under a 12-hour light-dark cycle. While the food was provided *ad libitum*, water was restricted during the training and experiment period (0.8 – 1.2 ml/day). Young adult mice were anesthetized with isoflurane (1 – 2 %) or with ketamine/xylazine. Lidocaine or bupivacaine was injected for pain relief and ketoprofen was injected for anti-inflammation. The skin and periosteum were removed, and a headpost was positioned on the skull. After 5 – 7 days of recovery, the mice were acclimated to the experiment environment under restricted water conditions.

For electrophysiological studies involving stimulation of thalamofrontal synapses, both wild-type C57BL/6J mice and transgenic mice generated by crossing PV-Cre (JAX: 008069) with Ai9 (JAX: 007909) were used. For ex vivo electrophysiological experiments, a Cre-dependent humanized channelrhodopsin (H134R) fused to enhanced yellow fluorescent protein (eYFP) was injected into the mediodorsal thalamus (MD) of PV-Cre x Ai9 (PV-tdTomato) mice. To enable Cre expression in MD neurons, AAV-hSyn-Cre (Addgene #105553) was co-injected. Stereotaxic coordinates for MD injection (relative to bregma) were as follows: anterior-posterior (AP) −1.20 to −1.58 mm, medial-lateral (ML) ±0.30 mm, and dorsal-ventral (DV) −3.10 mm. For in vivo electrophysiological experiments, AAV-CaMKllα-hChR2(E123T/T159C) was injected into the MD of wild type C57BL/6J mice to selectively evoke thalamofrontal activity during recordings.

All animal care and experimental procedures involving animals were approved by the Korea Brain Research Institute (KBRI) Institutional Animal Care and Use Committee or by the Johns Hopkins University Animal Care and Use Committee.

### Pharmacological inactivation

For local pharmacological inactivation, saline or 200 mM atropine (a muscarinic acetylcholine receptor (mAChR) antagonist; atropine sulfate salt monohydrate, Sigma-Aldrich) was infused through polyethylene tubing filled with mineral oil and connected with a 10μl-Nanofil syringe (WPI). Infusion was performed at the rate of 2 nl/s using an infusion pump (WPI) with total volume of 50 nl per injection site. After completion of the infusion, the cannula was left in place for several minutes to allow adequate diffusion of the infused solution. To minimize the effect of anesthesia, all infusions were performed while animals were awake. Behavioral tasks were conducted at least 1 hr after infusion to minimize potential behavioral alterations caused by stress associated with the injection procedure.

For systemic pharmacological antagonism, saline or atropine (10 mg/kg) was administered intraperitoneally (i.p.) at a volume of 0.02 ml, 30 min before the behavioral task.

### Electrophysiological recording during electrical and optogenetic stimulation of thalamofrontal synapses

#### ex vivo whole-cell recording

Mice were deeply anesthetized with CO_2_ and decapitated. Then brains were removed in ice-cold dissection solution (in mM): 25 Sodium bicarbonate (NaHCO_3_), 1.25 Sodium monophosphate (NaH_2_PO_4_), 25 D-Glucose, 2.5 KCl, 7 MgSO_4_·6H_2_O, 0.5 CaCl_2_, 110 Choline chloride, 11.61 (+) Sodiul-L-ascorbate, 3 Sodium pyruvate. Measured osmotic concentration was between 320 and 330 mOsm. 300-μm-thick acute brain slices containing dACC were prepared coronally by VT1200S Vibratome (Leica). The composition of artificial cerebrospinal fluid (aCSF) is (in mM): 119 NaCl, 2.5 KCl, 1 MgSO_4_·7H_2_O, 26 Sodium bicarbonate (NaHCO_3_), 1.25 Sodium monophosphate (NaH_2_PO_4_), 20 D-Glucose, 0.4 L-Ascorbic acid, 2 CaCl_2_, 2 Pyruvic acid. Measured osmotic concentration was between 305 and 310 mOsm. After 30 minutes of recovery in 32℃ aCSF, slices were transferred to room temperature. After PFC slices were prepared, slices containing the injection site were fixed for post hoc validation of the injection sites. Data from the mice with incorrectly targeted injections were excluded from data. The L2/3 pyramidal neurons in the dACC were recorded using whole-cell recording. Recordings were performed with patch pipettes (3.5 – 4 MΩ) filled with internal solution that consists of the following components (in mM): 20 KCl, 125 K-gluconate, 10 HEPES, 4 NaCl, 0.5 EGTA, 4 ATP, 0.3 Tris-GTP and 10 Phosphocreatine with pH adjusted to 7.20 with KOH. Measured osmotic concentration was between 307 and 314 mOsm. Recordings were performed at room temperature.

Unless otherwise indicated, optogenetic stimulation was delivered as 1-ms light pulses with a 470-nm laser source through an optic fiber positioned ∼1 mm away from the recorded neurons.

We measured the resting membrane potential in all neurons in current clamp mode immediately after the rupture of the neuronal membrane. Input resistance was determined by measuring voltage change in response to a small hyperpolarizing current pulse (5 pA, 50 ms) at resting potential. Rheobase current was acquired by 20 pA increasing step current injection and determined as the point at which the first action potential was evoked. Series resistance was monitored throughout each experiment and was not compensated. Cells with series resistance over 20 MΩ were excluded.

All solutions were kept saturated with 95% O_2_ and 5% CO_2_. Acute slices were continuously perfused with aCSF at room temperature. All data were sampled at 20 kHz by the EPC-10 amplifier (HEKA Elektronik) with Patchmaster software (HEKA Elektronik) and further analyzed by MATLAB (Math Works) and Igor Pro (WaveMetrics).

#### in vivo recording in anesthetized mice

For in vivo electrophysiological experiments, mice were anesthetized with ketamine (100 mg/kg, i.p.) and secured in a stereotaxic device (WPI, Sarasota, FL, USA). Anesthetic depth was checked by the toe-pinch reflex, and additional doses were given when necessary. A heating pad kept body temperature at 36.5 °C. After a scalp incision, a craniotomy was made above the recording site based on the atlas of Paxinos and Franklin (mPFC; AP: +1.7–2.0 mm from the bregma, ML: ±0.2–0.3 mm from the midline, DV: 0.8–1.2 mm below the dura), and the final position was adjusted until stable light-evoked responses appeared. A microscrew placed over the contralateral cortex served as the recording reference, and saline was applied to the exposed cortex to prevent drying.

Recordings were carried out 3–4 weeks after AAV-CaMKIIα-hChR2(E123T/T159C) was injected into the mediodorsal thalamus (MD). mPFC activity was acquired with a 16-channel optoelectrode that combines a silicon probe and a 200 μm fiber optic ferrule (A1×16-3mm-100-177-OA16LP; NeuroNexus, Ann Arbor, MI, USA), which was advanced slowly to ∼1,200 μm beneath the brain surface. MD axon terminals in the mPFC were activated with 2 ms pulses of 473 nm light delivered at 0.1 and 10 Hz. After baseline responses to both frequencies were obtained, carbamoylcholine chloride (CCh, 5 μM) was infused locally into the mPFC through a glass micropipette at 80 nL/min (UltraMicroPump and SYS-Micro4 Controller, WPI, Sarasota, FL, USA). Responses to 0.1 and 10 Hz photostimulation were measured again 5 min after CCh infusion, using identical stimulation parameters, to assess the effect of AChR activation.

Signals were amplified and digitized with a PZ5 NeuroDigitizer and an RZ5D BioAmp Processor (TDT, Alachua, FL, USA), sampled at 24,414 Hz per channel, and band-pass filtered from 0.3 to 5 kHz. Spikes were detected and sorted in Offline Sorter (Plexon Inc., Dallas, TX, USA) following our previous procedure (Yu et al., 2026). A detection threshold of four times the baseline noise level was applied, and artifacts were then removed. The detected spikes were grouped in principal component space by an automated valley-seeking algorithm with parameter optimization. Once outliers were removed, only units with significant cluster separation (MANOVA, p < 0.05) were accepted as single units. Their isolation was additionally confirmed from autocorrelograms (0.1 ms bins) computed in NeuroExplorer (Nex Technologies, Colorado Springs, CO, USA).

### Two-alternative forced-choice task

Mice were placed in a body restrainer and gently head-fixed in a custom-designed training chamber. Both auditory and visual 2AFC tasks consisted of three distinct epochs: sample, delay, and response.. During the sample period, mice were presented with either auditory or visual stimuli, depending on the task.

In the auditory task, behavioral procedures were controlled using an Arduino-based system. A 100-ms auditory tone of either 7 or 13 kHz was presented pseudo-randomly. In the visual task, visual stimulation and behavioral procedures were controlled using a Raspberry Pi, while hardware operation was managed by a custom circuit and an Arduino Mega. Vertical visual stimuli moving to “left” or “right” (“horizontal” or “vertical”) were presented pseudo-randomly on a monitor positioned in front of the animal for 4 s.

Following the sample epoch, mice were required to withhold licking during the delay period. The delay duration was initially set to 0 s and gradually increased up to 2 s as behavioral performance improved. After the delay, two lick ports were presented, and mice were required to report the presented stimulus by licking the corresponding left or right lick port during the response epoch. To minimize directional bias, cues of the same frequency were presented no more than three consecutive times.

Correct responses were rewarded with a small drop of water. In the auditory task, the ITI consisted of a fixed 2-s interval and an additional randomized interval drawn from a Gaussian distribution (mean = 1 s, range = 0-6 s). Following incorrect responses an additional 2-s penalty was added to the ITI. In the visual task, the ITI was fixed at 2 s and was extended to 4 s following incorrect responses, which were also signaled by a 250-ms auditory alarm tone (2 kHz).

Behavioral performance was evaluated using three measures: success rate, correct rate, and incorrect rate. Success rate was calculated as the proportion of correct trials among all trials, including omission trials. Correct and incorrect rates were calculated as the proportions of correct and incorrect responses, respectively, among completed trials, excluding omission trials.

### Fiber photometry recording in behaving mice

For fiber photometry, AAV9-hSyn-GRAB_ACh_3.0 (100 – 150 nl) ^35^ was bilaterally injected into the mPFC. Following viral injection, optical fiber ferrules (core diameter: 400 μm, NA: 0.22) were implanted bilaterally at the corresponding stereotaxic coordinates using a 30⁰ angled approach. Fiber photometry recordings were performed using a dual-color, multichannel fiber photometry system (R810, RWD Life Science, Shenzhen, China). ACh dynamics were monitored using the genetically encoded GRAB_ACh3.0 sensor with two excitation wavelengths: 470 nm for the ACh-sensitive fluorescence signal and 410 nm for the isosbestic reference signal. LED power measured at the tip of the optical patch cord was adjusted to approximately 20 μW for the 470 nm and 3 μW at the 410 nm to minimize photobleaching. Fluorescence signals were acquired at 30 frames s^-1^ using the scientific CMOS camera integrated into the system. Behavioral events were synchronized with the photometry recordings through TTL inputs to the acquisition system.

### In vivo recording in behaving mice

For in vivo extracellular recordings, a stainless-steel ground screw was implanted over the posterior cortex of either hemisphere, at a location distant from the recording site. A craniotomy was then made over the mPFC (AP: +1.94 mm from bregma; ML: ±0.4 mm from midline). The skull was thinned using a dental bur, and the remaining bone and dura were carefully removed using a tungsten needle. Between recording sessions, the craniotomy was covered with silicone elastomer (Kwik-Cast or Kwik-Sil) to protect the exposed brain surface and prevent tissue dehydration. Extracellular recordings were performed using Neuropixels 1.0 probes. Before insertion, the probe shank was cleaned with isopropyl alcohol and coated with the lipophilic fluorescent dye CM-DiI, DiD, or DiO dissolved in ethanol to enable subsequent histological reconstruction of the probe track. The probe ground was electrically connected to the implanted skull screw, and the probe was slowly lowered into the mPFC. After the probe reached the target recording depth (2.0 mm ventral to the brain surface), the exposed brain surface was covered with 1.5% agarose in artificial cerebrospinal fluid (aCSF) to maintain hydration and reduce brain movement. The probe was allowed to settle for approximately 10 min after insertion before behavioral recording began. Mice then performed the STM task while extracellular activity was recorded. Following each recording session, the probe was withdrawn and the craniotomy was resealed with silicone elastomer for subsequent recording sessions. Neural signals were acquired using SpikeGLX software (https://github.com/billkarsh/SpikeGLX)

## Data analysis

### Silicon probe recording preprocessing

#### Spike sorting and quality control

Extracellular electrophysiological recordings obtained via Neuropixels probes were preprocessed and spike-sorted to isolate single unit activity. To isolate high-frequency spiking activity, the extracellular AP signals were high-pass filtered using an acausal filter with a cutoff frequency of 300 Hz. Common-mode noise and channel cross talk were reduced using common-average referencing (CAR). Automated spike sorting was performed on the preprocessed continuous binary data using Kilosort2.5 via the open-source pipeline (https://github.com/jenniferColonell/ecephys_spike_sorting). Template matching, drift correction, and cluster identification were executed with default parameter sets, using a threshold behavior dictionary optimized for cortical recordings.

Following the automated sorting, all identified clusters were subjected to rigorous manual curation using Phy2 and classified as single-unit activity (‘good’), multi-unit activity (mua), or noise based on waveform characteristics, refractory-period violations, and cluster quality.

Units classified as ‘good’ were further subjected to quantitative quality-control criteria. Units with an inter-spike interval (ISI) violation metric >0.5 or an amplitude cutoff >0.1 were excluded from subsequent analyses. The amplitude-cutoff criterion was used to exclude units with a substantial estimated fraction of missed spikes. After curation and quality filtering, the final dataset comprised 430, 272, and 253 single units in the control, atropine, and saline conditions, respectively. Only these curated single units were included in subsequent neurophysiological analyses.

#### Peri-stimulus time histogram (PSTH)

After aligning the neural activity according to the timing of the stimulus onset, the PSTH was calculated as the number of spikes in 1 ms bins, followed by smoothing with a 100 ms boxcar filter. The neural firing activity obtained from the inter-trial interval (ITI), 2 sec before the start of each trial, was used as the baseline for normalization. Each PSTH was z-score normalized by subtracting the mean firing rate of the baseline and dividing by the standard deviation of the firing rate of the baseline.

### Single unit analysis

#### Directional selectivity of single units

Trial types were classified according to task epochs as follows: stimulus (the period in which either a high tone or low tone auditory cue was presented), delay (the interval during which the animal deliberated whether to lick left or lick right), and response (the period in which a correct or incorrect response was executed). We computed neural selectivity separately for each trial type. To calculate selectivity, the trial types were combined into four variables (stimulus left and choice left 〈*LL*〉; stimulus left and choice right 〈*LR*〉; stimulus right and choice right 〈*RR*〉; stimulus right and choice left 〈*RL*〉), and then converted into two corresponding groups for each trial type as follows ^36^:

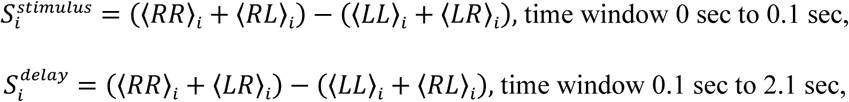

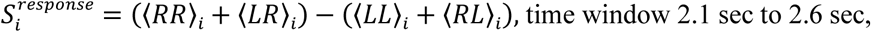

where *i* denotes the neural index, and each variable was calculated by averaging the baseline-subtracted PSTH within the corresponding time window. Neural selectivity was then examined as a significant difference between the mean PSTH of the two groups using the Mann-Whitney U test.

#### ROC curve analysis

To investigate whether each neuron encoded direction, we computed and analyzed ROC curves using MATLAB’s *perfcurve* function on trial-averaged PSTH obtained after applying a 100 ms boxcar filter and baseline subtraction. For this analysis, only the task period of the PSTH, excluding the baseline period, was used. The positive class was defined as the direction (left licking PSTH vs. right licking PSTH) associated with the higher mean firing rate. ROC curves were computed separately for correct licking trials and incorrect licking trials. For bootstrapping, 80% of trials were randomly sampled for each unit to compute the trial-averaged PSTH, and this procedure was repeated 100 times to generate ROC curves. For each neuron, the decoding performance was quantified as the mean auROC value, obtained by averaging the areas under bootstrapped ROC curves. From this, we defined a sensitivity index (*SI*) as follows: *SI* = |auROC − 0.5| × 2. Both the auROC and the sensitivity index were binned from 0 to 1 with a step size of 0.01, and histograms were constructed using MATLAB’s *histcounts* function.

To calculate a data-driven chance level, we performed shuffling in two ways. First, trial-level shuffling was performed by pooling the training trials from the two compared directions and randomly repartitioning them into two pseudo-direction groups while preserving the original number of trials. The trial-averaged PSTH was then computed from these shuffled trial groups, and ROC curves and auROC values were obtained using the same procedure as in the original analysis above. This procedure was applied separately to correct trials and incorrect trials, thereby breaking the relationship between trial identity and licking direction while preserving the trial-level firing rate structure. Second, label shuffling was performed at the trial-averaged PSTH level. In this analysis, trials were first averaged according to their original direction labels to generate the true direction-specific PSTH. The direction labels provided to MATLAB’s *perfcurve* function were then randomly permuted while preserving the number of labels assigned to each direction. The positive class was again defined as the shuffled label group with the higher mean firing rate, using the same criterion as in the original auROC analysis. For both shuffling procedures, the same smoothing, baseline subtraction, task-period selection, 80% trial sampling, and 100 bootstrap repetitions were used as in the original analysis. The resulting shuffled auROC values were used to estimate the chance-level null distribution of decoding performance.

#### Distribution of differences in mean firing rates across epochs

To quantify differences in neuronal activity between RR and LL trials across task epochs, trial-averaged PSTHs were calculated separately for each trial type using 1 ms bins without temporal smoothing. For each neuron, each PSTH was z-score normalized by subtracting its mean baseline firing rate and dividing by its baseline standard deviation. Normalized firing rates were then averaged within delay epochs: early delay (0.1–1.1 s) and late delay (1.1–2.1 s). For each neuron and epoch, the absolute difference between the normalized mean firing rates for RR and LL trials was calculated.

To compare the two conditions with equal sample sizes, we drew 100 units at random with replacement from each pooled population. We then calculated the proportion of sampled units classified as direction-selective in each epoch, using the same sampled units for both epochs. This procedure was repeated 100 times. Each repetition yielded one proportion of direction-selective units per condition and epoch. For each epoch, the 100 proportions from the pre-injection condition were compared with those from the atropine condition using a two-sided Mann–Whitney U test, with Bonferroni correction across epochs. The relative reduction in the proportion of selective units was calculated for each repetition as (P_be*F*ore_ − P_atropine_)/P_be*F*ore_. Early- and late-delay reductions were compared using a two-sided Mann–Whitney U test, excluding repetitions in which the pre-injection proportion was zero for the corresponding epoch.

### Neural population analysis

To examine the changes in sustainability of working memory, we used three complementary methods for neural population analysis: (1) Pearson correlation analysis, (2) linear discriminant analysis, and (3) coding direction analysis.

#### Pearson correlation analysis

We first analyzed the neural population activity in its native space, in which the dimensionality was not reduced. To achieve this purpose, we calculated the Pearson correlation coefficient ^37^ between trial-averaged population vectors at time points *t*_1_ and *t*_2_:

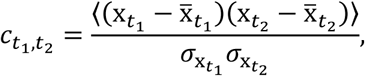

where x*_t_*_1_ and x*_t_*_2_ are the trial-averaged PSTH at time points *t*_1_ and *t*_2_, respectively. To compare the variability of correlation coefficients across groups, we defined the diagonal-averaged correlation as the average of the diagonal elements in the matrix of Pearson correlation coefficients, as follows:

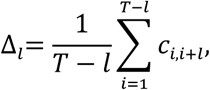

where *T* is the total number of time points for the neural population activity. We assumed that the diagonal-averaged correlation decays exponentially with lag *l*: Δ(*l*) ≈ *A* ⋅ *e*^−*λλ*⋅^*^l^*. By performing a linear regression of ln Δ(*l*) = ln *A* − *λ* ⋅ *l* against lag *l*, we estimated the decay rate *λ*. The time constant *τ* was then computed as *τ* = ln 2 /(−*λ*). This time constant represents how rapidly the Pearson correlation coefficient decays over time. To compensate for differences in the number of neurons across groups, we randomly sampled 100 neurons from each group and repeated this procedure 100 times. For each sampled population, we fitted the correlation decay curve and obtained one decay constant. Pairwise comparisons were performed on these decay constants using two-sided Mann–Whitney U tests with Bonferroni correction for the four comparisons.

We also calculated Pearson correlations between late-delay and early-delay population activity and between late-delay and response-period population activity. Within each condition, both correlations were calculated using the same 100 neurons in each repetition. Correlations were calculated separately for correct right- and left-licking trials and then averaged across the two directions. This yielded one pair of correlation coefficients per repetition. The two coefficients were compared within each condition using a Wilcoxon signed-rank test across 100 paired repetitions.

#### Linear discriminant analysis

To evaluate how much neural population activity contained trial type- and epoch-related information, we applied LDA. For both correct and incorrect trials, baseline-subtracted single-trial firing rates were used to construct pseudo-population activity. For decoding based on activity averaged over the task period, original trials within each session and trial type were randomly divided into non-overlapping training and test sets before pseudo-population samples were constructed, with 80% of trials assigned to the training set and the remaining 20% assigned to the test set. LDA classifiers were trained separately for correct and incorrect trials. For correct trials, classifiers were trained to discriminate right-licking correct trials from left-licking correct trials. For incorrect trials, classifiers were trained to discriminate right-licking incorrect trials from left-licking incorrect trials. Classification of population activity averaged over the task period was performed using MATLAB’s *fitcdiscr* function with a pseudo-linear discriminant model and empirical class priors. Decoding performance was quantified from classifier scores by computing ROC curves using MATLAB’s *perfcurve* function, with the right-licking condition defined as the positive class. Each held-out pseudo-population sample contributed one right-licking classifier score, labeled by the actual licking direction. In the single-neuron ROC analysis, each score supplied to *perfcurve* was the firing rate in one time bin of a trial-averaged PSTH, labeled by the corresponding licking direction.

We performed two sets of population decoding analyses. First, to determine whether licking direction could be decoded more effectively from neural population activity than from single unit activity, we performed LDA using the entire neural population recorded in the before group. In this analysis, pseudo-population activity was constructed by concatenating all recorded units across sessions. Each neuron’s activity was smoothed, baseline-subtracted, and averaged over the task period, resulting in one activity value per neuron for each pseudo-population sample. This procedure generated 100 pseudo-population samples per trial type for each repetition, with 80 samples used for training and 20 samples used for testing. The analysis was repeated 100 times to obtain a robust estimate of decoding performance for correct and incorrect trials.

Second, to compare decoding performance across groups while controlling for differences in neuron and trial counts, we performed time-resolved LDA analyses using single-trial activity with matched neuron and trial counts across groups. Two population configurations were examined: pseudo-populations sampled from the full unit pool and pseudo-populations sampled after excluding units exhibiting statistically significant directional selectivity during the response epoch. For each group and configuration, 100 units were randomly sampled without replacement, and the complete sampling and decoding procedure was repeated 100 times. For each selected unit, trials were independently sampled without replacement, with trial counts matched across units, trial types, and groups, and combined to construct pseudo-population trials. Each decoding feature was calculated as the baseline-subtracted mean firing rate within a causal 200 ms sliding window, with a step size of 50 ms. LDA was fitted separately at each time point and evaluated using 5-fold cross-validation across pseudo-population trials. For each fold, auROC was calculated from the ranks of the held-out discriminant scores, with right licking as the positive class. The resulting auROC values were averaged across folds.

For comparisons of decoding performance, the observations were auROC values obtained from 100 resampling repetitions per condition. The repetitions sampled the same underlying recordings and could include the same neurons across repetitions. To compare decoding across conditions using activity averaged over the task period, we sampled 100 neurons per condition in each repetition and obtained one auROC value for correct trials. We compared these values across the pre-injection, saline, and atropine conditions using two-sided Mann–Whitney U tests with Bonferroni correction for the three pairwise comparisons. In the time-resolved analysis, each repetition contributed one fold-averaged auROC value per time point for each condition, population configuration, and trial outcome.

#### Coding direction analysis

To visualize direction-selective population activity, we performed a coding direction analysis ^5,38^. For a neural population of *n* neurons, we sought an *n* × 1 basis vector that would maximally discriminate the two trial types during the delay epoch. At each time point *t*, we defined the direction-selective population vector *r_t_* = *r_lick_*_−*rrig*ℎ*t,t*_ − *r_lick_*_−*left,t*_, where *r_lick_*_−*rrig*ℎ*t,t*_ and *r_lick_*_−*left,t*_ are *n* × 1 vectors of baseline-subtracted PSTH for individual neurons averaged across correct right- and left-lick trials, respectively. Because activity during the delay epoch exhibits sustained, relay-like dynamics in the brain regions under consideration, we computed the basis vector of coding direction *CD_delay_ = 1/T_delay_Σ_tT_* followed by normalization with its norm. Similarly, *CD_stim_* and *CD_rresponse_* were obtained by averaging *r_t_* over all stimulus and response time points, respectively. To derive the go-direction vector *D_go_*, we first computed the trial-type-independent mean activity (*r_lick_*_−*rrig*ℎ*t,t*_ + *r_lick_*_−*left,t*_)/2, then formed the difference between the mean firing rate immediately after the response execution and that immediately before response onset, followed by normalization. We defined the ramping direction *D_rramp_* as the unit vector that maximally separated the average population activity just prior to trial onset from that immediately before response execution. All direction vectors were orthogonalized using the Gram-Schmidt process. This coding direction analysis was performed using 5-fold cross-validation across trials for each neuron. Specifically, 80% of trials were used to compute trial-averaged PSTH, from which all coding direction vectors were derived. The remaining 20% of trials were used to compute another trial-averaged PSTH, which was then projected onto each direction-selective basis to quantify the coding direction. Because the neural response subspace could differ across neurons, we further normalized the coding direction estimates by z-scoring them relative to the mean and variance obtained from coding directions computed using randomly shuffled PSTH within each group. Specifically, we generated surrogate datasets by randomly permuting the temporal order of the training and test PSTH within each fold. For each bootstrap iteration, the same temporal permutation was applied to all trial types, a shuffled coding direction basis was recomputed from the shuffled training PSTH, and the correspondingly shuffled test PSTH were projected onto the resulting basis. This procedure was repeated 100 times per fold. For each fold and time bin, the projection obtained from the unshuffled data was z-scored relative to the mean and standard deviation of the shuffled projections.

### Clustering of individual neuron response profiles

#### ePAIRS test for clustering of response profiles

We first assessed whether neuronal response profiles showed evidence of clustering by applying the elliptical projection angle index of response similarity (ePAIRS) test ^39^. The input comprised an *n* × *t* matrix of neural population firing rates, where each row represented the PSTH of an individual neuron and the ‘lick left’ and ‘lick right’ trials were concatenated. All responses were baseline-subtracted and magnitude normalized. We used PCA to reduce the temporal dimensionality from *t* to 37 that account for 90% of the activity variance over time ^39^. To estimate the response similarity, we computed the average vector angle between each neuron and its *k* nearest neighbors in the reduced dimension as follows: *α_i_ = 1/K.Σ_jNi_θ^(i)^_j_*, where α is the average vector angle between the neuron *i* and its *k* nearest neighbors. The subset *N_i_* denotes the indices of neurons which are the *k* nearest neighbors of the neuron *i* based on the Euclidean distance. The angle between the response vectors of neurons *i* and *jj* is estimated as follows: *θ^(i)^_j_ = arccos((x_i_.x_j_)/(‖x_i_‖.‖x_j_‖)*. We obtained a distribution of averaged angles across the neural population.

To assess whether response profiles were clustered, we generated null distributions that possessed the same statistical properties as the dimensionally reduced neural population activity but exhibited no clustering. We drew *n* samples from a 37-dimensional multivariate Gaussian distribution with zero mean and equal variance in each dimension of the neural data ^38,39^ using the MATLAB function *mvrnd()*. To test the statistical difference between the null distribution of the simulated dataset and the empirical distribution, we computed the averaged vector angles from the simulation and repeated 10,000 times. The significance *p* value was calculated as the proportion of cases in which the median of the empirical distribution exceeded the median of the null distribution. We tested a range of *k* values (1–10) and found that, for each *k*, none of the 10,000 simulated null medians was smaller than the empirical median.

#### t-SNE embeddings and density peak clustering of response profiles

We used the t-distributed stochastic neighbor embedding (t-SNE) ^40^ method to reduce the dimensionality of the *n* × 2*t* matrix of PSTH, formed by concatenating the PSTH from correct right-and left-licking trials along the time dimension, from 2*t* to 3. This approach yielded more consistent PSTH within the clusters than those obtained using the top principal component approach ^38^. We used perplexity of 50, 50 principal components, and cosine distance ^38^ using the MATLAB function *tnse()*. The embedding was run ten times, and we selected the result with the lowest Kullback-Leibler divergence. Embeddings were performed separately for the before and atropine groups. We then performed the density peak clustering ^41^ in the three-dimensional subspaces obtained from the t-SNE method. The key hyperparameter of density peak clustering, the cutoff distance *d_c_*, was set as in the original paper ^41^ to the value corresponding to the 2% of the pairwise distance distribution obtained using a Gaussian kernel. Density peak clustering then requires manual selection of cluster centers based on the local density *ρ* and distance to higher-density neighbor *δ*. We employed the gamma strategy ^42,43^ to improve consistency and reproducibility in the number of clusters and center selection by defining the decision value as *γ_i_* = *ρ_i_* ⋅ *δ_i_*, where *i* indexes each unit. We then sorted all *γ_i_* values in descending order, producing a curve with an exponentially decreasing shape, and fitted two lines to this curve. By computing the sum of absolute errors for both regressions, we identified the knee point as the position at which this error sum is minimized ^42,43^. The *ρ_i_* and *δ_i_* values corresponding to that knee point were chosen as a compromise between under- and over-clustering and used to perform density peak clustering. Clusters with similar PSTH were merged, resulting in three clusters each for the before and atropine groups.

## RESULTS

### Frequency-dependent gating of MD–dmPFC transmission

Given the proposed role of the TCT activity loop in sustaining memory-relevant activity in the PFC, we asked whether short-term synaptic depression in the MD to PFC pathway could limit the fidelity with which sustained thalamic activity is transmitted to the PFC. Although short-term synaptic depression is a hallmark of first-order thalamocortical synapses in sensory systems ^15,34,44^ and MD-to-PFC synapses ^45,46^, the extent to which this property constrains sustained thalamofrontal transmission remains unclear.

To examine how short-term depression influences thalamofrontal transmission, we expressed hChR2-eYFP in MD neurons and optogenetically stimulated MD axons while recording from L2/3 pyramidal neurons in acute mPFC slices (Fig. 1A). MD axons densely innervated the mPFC, with strongest projections to L2/3, consistent with previous reports (Fig. 1A, bottom) ^46–48^. Low-frequency MD stimulation reliably evoked action potentials in L2/3 pyramidal neurons, whereas repetitive high-frequency stimulation progressively reduced spike probability, with transmission deteriorating markedly at frequencies ≥5 Hz (Fig. 1B). These findings demonstrate that frequency-dependent short-term depression imposes a substantial constraint on the reliable transmission of sustained thalamic activity to the mPFC.

**Figure 1.**
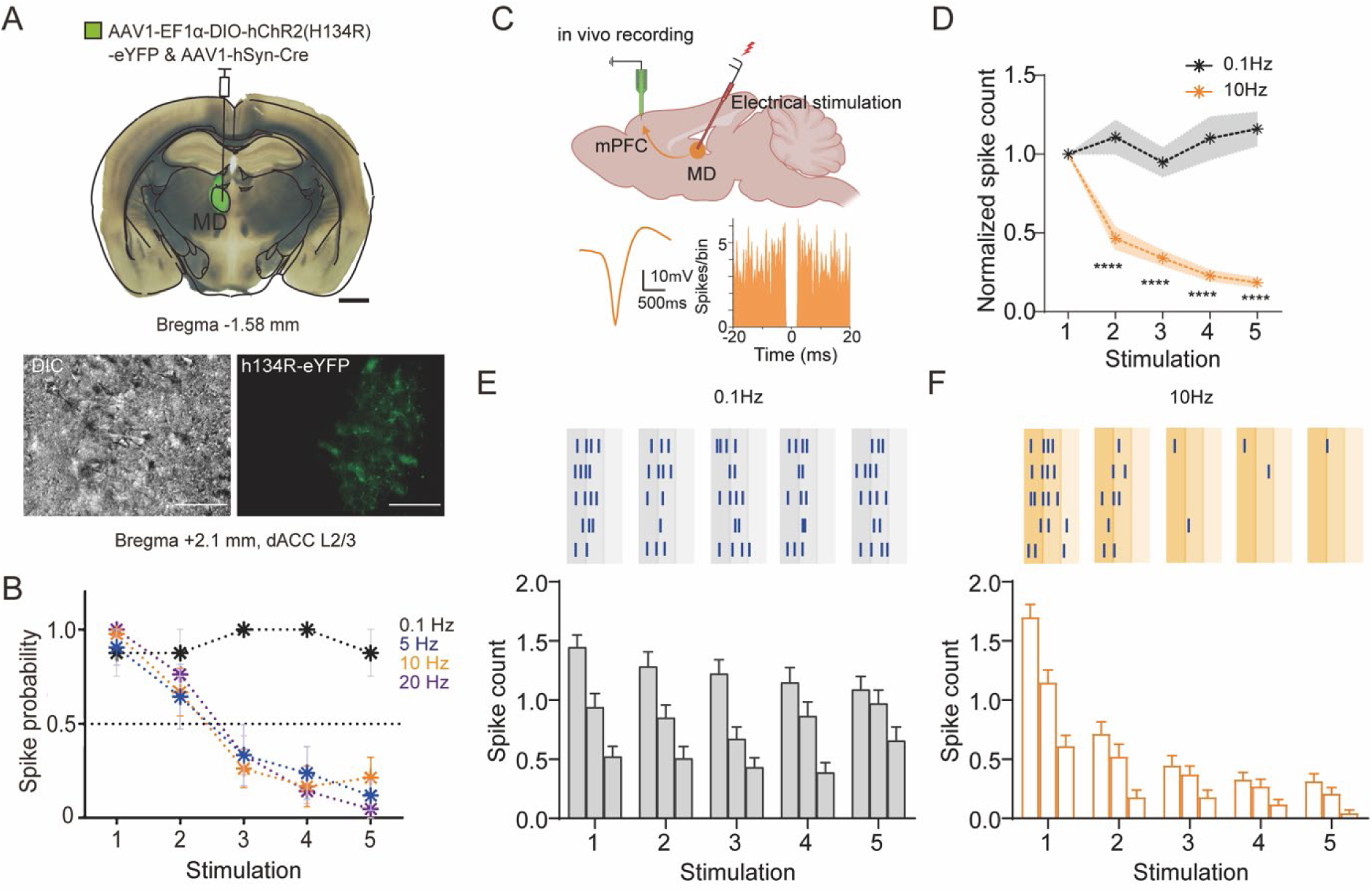
Frequency-dependent depression limits thalamofrontal transmission. **(A)** Top, representative coronal section showing ChR2-eYFP expression following viral injection into the MD. Scale bar: 1 mm. Bottom, representative differential interference contrast (DIC) and fluorescence images showing ChR2-eYFP expressing axon terminals in layer 2/3 of the dorsal anterior cingulate cortex (dACC), Scale bar: 10 μm. **(B)** Spike probability of L2/3 pyramidal neurons during repetitive optogenetic stimulation of MD axons at 0.1, 5, 10, and 20 Hz *ex vivo*. **(C)** Top, schematic of electrical stimulation of the MD and simultaneous extracellular recording in the mPFC *in vivo*. Bottom, representative waveform of an isolated unit and autocorrelogram. **(D)** Normalized spike counts evoked by five consecutive stimuli delivered at 0.1 Hz (black) or 10 Hz (orange) (\*\*\*\**p*<0.00001). **(E-F)** Representative spike raster plots (top) and corresponding spike counts measured in sequential 10-ms bins (bottom) during MD stimulation at (E) 0.1 Hz or (F) 10 Hz in vivo. Data are shown for optogenetic stimulation of thalamofrontal axons and electrical stimulation of the MD.

Because frequency-dependent reductions in spike probability could reflect ChR2 inactivation, we first examined the reliability of light-evoked spiking in MD neurons. Spike-generation reliability remained comparable across stimulation frequencies at a given pulse duration (Extended Data Fig. 1A and Table 1), arguing against ChR2 inactivation as the primary cause of the observed depression.

Because both physiological extracellular Ca²⁺ concentrations and ongoing network activity can substantially reduce the magnitude of short-term synaptic depression observed in vitro ^49,50^, we next asked whether repetitive MD activity is similarly filtered in the intact brain. Extracellular recordings from the mPFC during electrical stimulation of the MD revealed pronounced frequency-dependent attenuation of cortical responses in vivo, closely paralleling the depression observed *ex vivo* (Fig. 1C-F). These findings demonstrate that the frequency-dependent bottleneck is preserved in the intact thalamofrontal pathway.

To determine whether the observed depression arose at monosynaptic thalamofrontal synapses, we isolated monosynaptic ChR2-evoked responses using TTX and 4-AP ^51–53^. Under these conditions, optically evoked EPSCs remained robustly depressing (Extended Data Fig. 1B-E), demonstrating that frequency-dependent depression is an intrinsic property of monosynaptic thalamofrontal transmission and does not require recurrent cortical circuitry. Together, these findings demonstrate that thalamofrontal synapses impose a frequency-dependent bottleneck on sustained thalamocortical communication, reliably transmitting low-frequency inputs while progressively attenuating repetitive activity.

### STM task and acetylcholine

Given that MD→PFC synapses exhibit strong short-term depression, yet disruption of this pathway during the delay period impairs STM performance ^10^, we reasoned that additional mechanisms must preserve thalamofrontal transmission during behavior. Because cholinergic signaling has been implicated in attention and STM ^24^, we hypothesized that ACh dynamically supports thalamofrontal information transfer by compensating for the progressive loss of synaptic efficacy during repetitive activity.

To test this hypothesis, we monitored extracellular ACh dynamics in the mPFC using the genetically encoded ACh sensor, GRAB_ACh_3.0 ^35^ and fiber photometry while head-fixed mice performed an auditory delayed-response task (Fig. 2A). In each trial, mice were presented with a 100-ms auditory cue (7 or 13 kHz), maintained cue information across a 2-s delay, and subsequently reported cue identity by licking the corresponding response port for reward (Fig. 2B).

**Figure 2.**
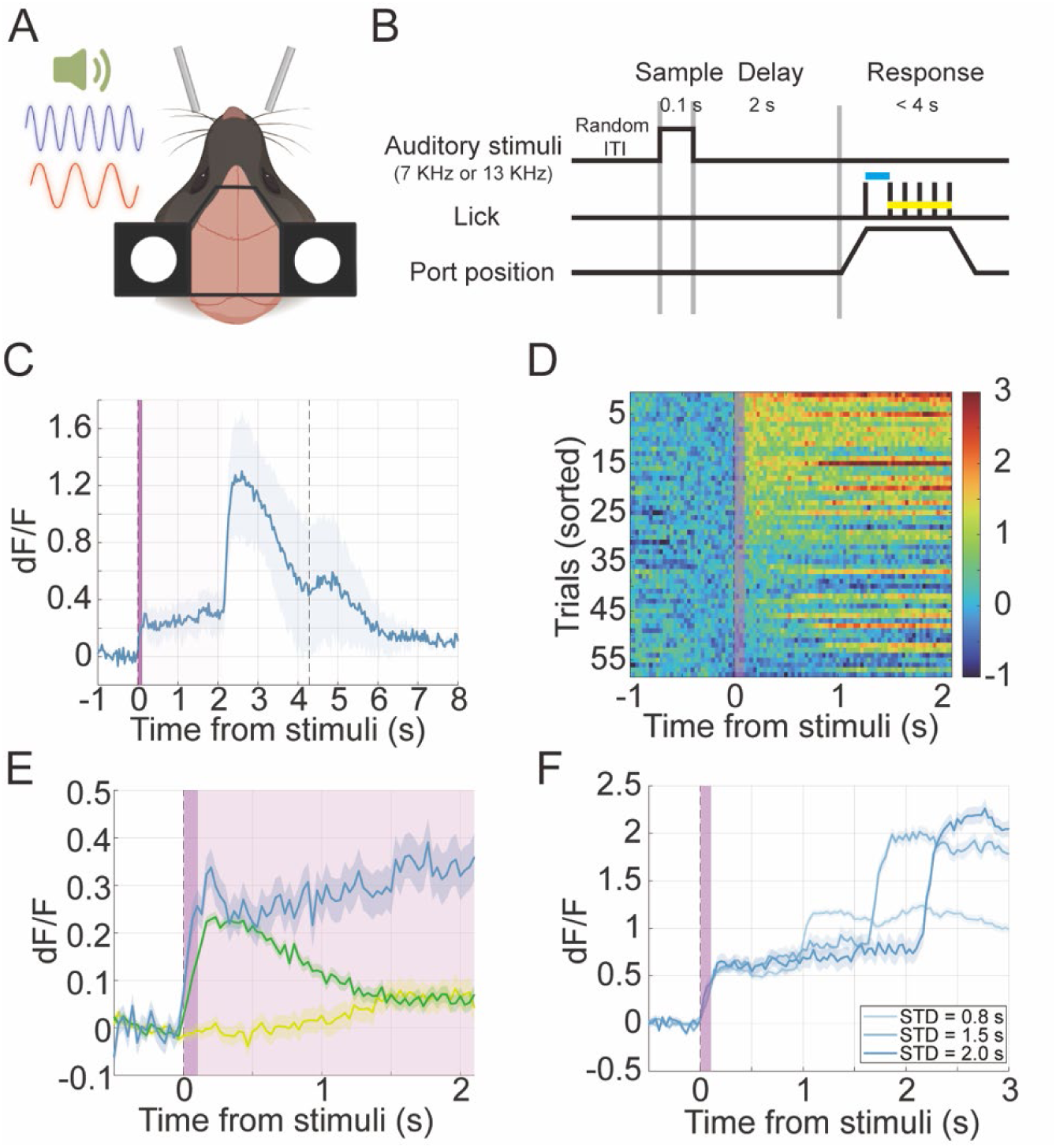
Prefrontal acetylcholine dynamics during an auditory STM task. **(A)** Schematic of the head-fixed behavioral setup. **(B)** Structure of the auditory delayed-response paradigm. A 100-ms auditory cue (7 or 13 kHz) was followed by a 2-s delay and subsequent presentation of the lick ports for behavioral response. **(C)** Representative ACh signal (dF/F) aligned to auditory stimulus onset (mean ± SEM). The purple bar indicates cue period (100 ms) and the light purple shaded region indicates the 2-s delay period. The vertical dashed line denotes lick-port retraction. **(D)** Single-trial heatmap of ACh signals from a representative session aligned to cue onset. Trials are sorted by the latency of the cue-evoked increase in dF/F. Color indicates normalized ACh signal magnitude. **(E)** ACh dynamics in well-trained mice (blue) and task-naïve mice presented with (green) and without (yellow) an auditory cue. **(F)** ACh dynamics during trials with different delay durations, showing that the duration of the elevated ACh signal scales with the imposed delay interval, while a second elevation occurs during reward consumption.

Extracellular ACh increased rapidly following cue presentation, remained elevated throughout the delay period, and increased further during reward consumption (Fig. 2C, D). To determine whether the sustained cholinergic signal reflected a prolonged sensory response, we compared ACh dynamics across training states and task conditions. In task-naïve mice, no-cue trials produced no detectable increase in the ACh signal (Fig. 2E and Extended Data Fig. 2A, yellow), whereas cue presentation evoked a rapid transient increase in both task-naïve and well-trained mice (Fig. 2E and Extended Data Fig. 2A, blue and green). Importantly, however, the cue-evoked signal returned toward baseline in naïve animals, whereas it remained elevated throughout the delay during task performance. We next varied the delay duration to determine whether this sustained component tracked the maintenance interval. ACh remained elevated during the tested delay intervals. (Fig. 2F and Extended Data Fig. 2B,C). These findings indicate that a transient cholinergic response can be triggered by the sensory cue itself, whereas sustained ACh elevation is associated with task engagement and tracks the duration over which information must be maintained. We next asked whether delay-period cholinergic activity predicted subsequent behavioral performance. ACh increased rapidly after cue presentation and remained elevated throughout the delay in both correct and incorrect trials (Extended Data Fig. 2D,E). Following reward consumption in correct trials or the first incorrect lick in unrewarded trials, the ACh signal decayed toward baseline within approximately 1 s (Extended Data Fig. 2). The rapid return toward baseline after completion of the response epoch further indicates that the elevated cholinergic state is temporally confined to ongoing task engagement.

### Cholinergic regulation of thalamofrontal transmission and dmPFC neuronal excitability

Having observed sustained ACh elevation during the memory delay, we next asked whether cholinergic signaling could enhance the ability of repetitive thalamic input to drive prefrontal neurons. We applied the non-selective cholinergic receptor agonist carbachol (CCh, 5 μM) during whole-cell recordings from dmPFC pyramidal neurons in acute brain slices. CCh significantly increased spike probability during high-frequency stimulation of MD axons (Fig. 3A–B). A comparable enhancement of cortical responses was observed in anesthetized mice during repetitive electrical MD stimulation (Extended Data Fig. 3A–B).

**Figure 3.**
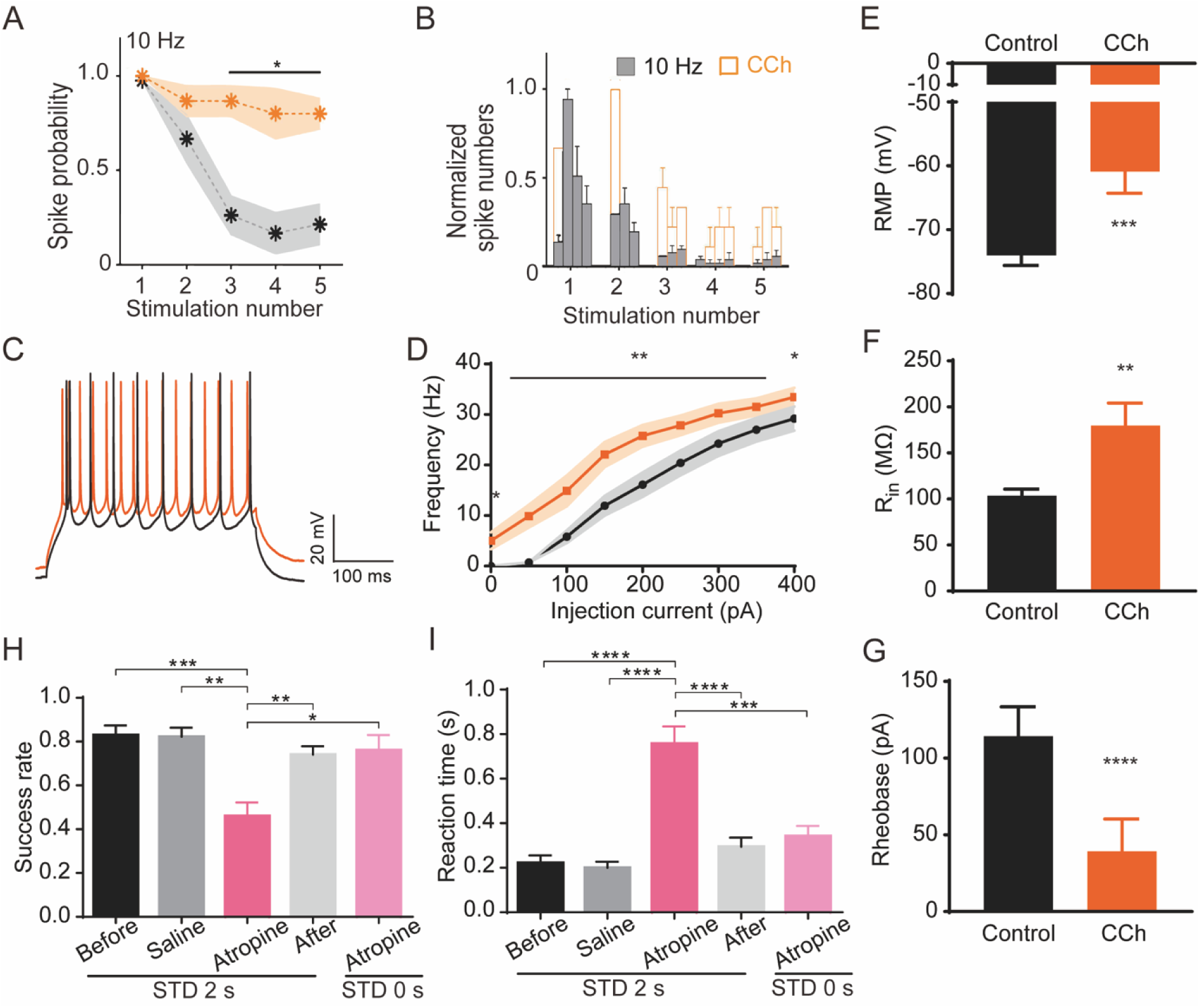
Cholinergic modulation of thalamofrontal transmission, prefrontal excitability, and STM performance. **(A)** Spike probability of dmPFC neurons during five consecutive optogenetic stimuli of MD axons at 10 Hz in acute brain slices, under control conditions (black) and during CCh application (orange). Asterisks indicate the mean and shaded areas indicate SEM (\**p*<0.05, paired t-test). **(B)** Normalized spike responses recorded ex vivo from the dmPFC during repetitive MD stimulation at 10 Hz under control conditions (black) and following CCh application (orange) (bin width 10 ms). **(C)** Representative membrane potential traces during current injection in control (black) and CCh (orange). **(D)** Firing frequency as a function of injected current. **(E-G)** Summary of intrinsic excitability changes showing (E) depolarized resting membrane potential, (F) increased input resistance, and (G) reduced rheobase current following CCh bath application. **(H-I)** Selective impairment of STM performance following mAChR antagonism, showing (H) decreased success rate and (I) increased reaction time without affecting the delay-free discrimination task (n=4, 19 sessions, one-way ANOVA).

We next asked whether cholinergic signaling alleviates the frequency-dependent bottleneck at thalamofrontal synapses by reducing presynaptic release probability ^54–55^. Such a reduction would be expected to attenuate short-term synaptic depression and thereby improve the transmission of sustained MD activity to the dmPFC. However, cholinergic receptor activation did not alter short-term synaptic plasticity at thalamofrontal synapses (Extended Data Fig. 3C-D). Instead, CCh depolarized dmPFC pyramidal neurons, increased their input resistance, and reduced their rheobase, consistent with enhanced intrinsic excitability (Fig. 3C-G) ^56–58^. By contrast, nicotine produced only a small depolarization but did not alter input resistance or rheobase (Extended Data Fig. 3E-I), indicating that the increase in intrinsic excitability is predominantly mediated by muscarinic receptors.

### Cholinergic control of prefrontal function during memory-guided behavior

Having established the cellular mechanism by which cholinergic signaling enhances thalamofrontal transmission, we next asked whether endogenous muscarinic signaling is required for STM performance in behaving animals. Systemic administration of atropine reversibly reduced overall task accuracy to chance levels, whereas saline administration had no effect (Fig. 3H, including omissions). To determine whether this impairment reflected deficits in sensory discrimination or motor execution rather than memory maintenance, we repeated the task under a zero-delay condition. Performance remained largely intact, indicating that atropine did not impair cue discrimination or directional licking per se (Fig. 3H). Atropine nevertheless increased reaction time and the proportion of omission trials (Fig. 3I; Extended Data Fig. 4A), raising the possibility that reduced task engagement contributed to the behavioral deficit. However, when the analysis was restricted to completed trials, atropine still significantly reduced response accuracy, with fewer correct and more incorrect choices than during saline sessions (Extended Data Fig. 4B-C, excluding omissions). Thus, the impairment in STM performance could not be explained solely by reduced task engagement. Rather, atropine also impaired behavioral performance on trials in which mice initiated a response, consistent with a deficit in maintaining or utilizing mnemonic information.

Together, these findings support a model in which endogenous muscarinic signaling enhances effective thalamofrontal transmission and thereby contributes to STM performance.

Given that atropine impaired choice accuracy even when omission trials were excluded, we focused our subsequent analyses on completed trials and compared neural activity between correct and incorrect choices. This approach allowed us to examine the neural correlates of behavioral accuracy while minimizing potential confounds associated with trial omissions and response initiation.

### Sequential dynamics and population-level directional encoding in dmPFC

We next characterized the organization of task-related dmPFC activity during STM using extracellular recordings with Neuropixels probes in behaving mice (Fig. 4A). During task performance, dmPFC neurons exhibited diverse task-related response profiles spanning the cue, delay, and response periods (Fig. 4B). Together, these neurons formed a temporally organized population representation of the task, consistent with previous reports of sequential activity in the PFC during STM ^10^.

**Figure 4.**
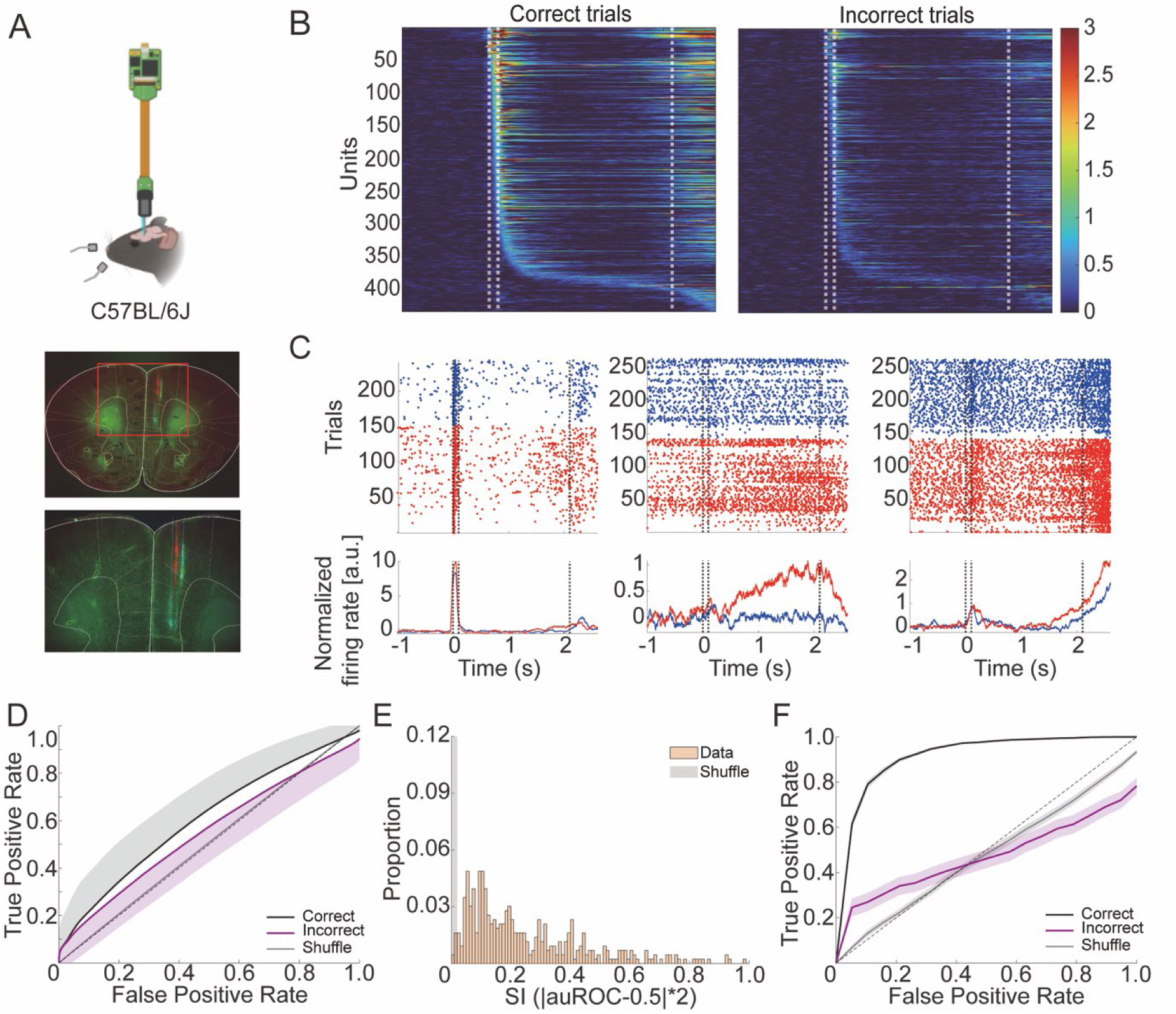
Sequential activity and directional selectivity of dmPFC neurons during the STM task. **(A)** Schematic of in vivo high-density extracellular recording in head-fixed mouse (top) and representative coronal section showing the electrode tracks in the dmPFC. **(B)** Sequential activity of isolated dmPFC units during correct (left) and incorrect (right) trials. Units are ordered according to the latency of peak firing during correct trials. **(C)** Representative raster plots (top) and normalized peri-stimulus time histograms (PSTHs, bottom) from three units exhibiting distinct temporal profiles. Blue and red indicate left- and right-cued trials, respectively. **(D)** Single-unit receiver operating characteristic (ROC) analysis of directional selectivity during correct (black) and incorrect (purple) trials. Curves show the population average across individual neurons; shuffled trial labels are shown in gray. **(E)** Distribution of single unit selectivity indices (SI = |auROC - 0.5| x 2) for recorded mPFC units (data) and trial-shuffled controls. **(F)** Population-level decoding of trial direction using a cross-validated linear discriminant analysis (LDA) classifier. ROC curves show decoder performance for correct (black) and incorrect (purple) trials relative to shuffled controls (gray).

Further, a subset of neurons exhibited clear directional selectivity, with differential activity depending on the upcoming lick direction (Fig. 4C, red vs. blue). However, the average direction selectivity across the population was only moderate, as quantified by receiver operating characteristic (ROC) analysis (Fig. 4D). We therefore asked whether directional information was preferentially represented by a particular temporal response class. To address this question, we embedded individual neuronal activity into a low-dimensional space using t-distributed stochastic neighbor embedding (t-SNE), followed by density peak clustering. This unsupervised analysis identified three major response classes that closely matched our manual classification: cue-responsive neurons, neurons exhibiting sustained delay activity, and neurons displaying ramping activity preceding the response (Extended Data Fig. 5A). ROC analysis performed separately within each response class revealed comparable levels of directional selectivity across all three groups (Extended Data Fig. 5B), indicating that directional information is broadly distributed across temporally distinct neuronal populations rather than being confined to a specialized response class. Nevertheless, the distribution of selectivity indices (SI; Methods) was significantly right-shifted relative to chance, revealing a small subset of strongly direction-selective neurons (Fig. 4E and Extended Data Fig. 5C-D).

Together, these findings indicated that directional information is broadly distributed across the dmPFC, with individual neurons exhibiting only modest directional selectivity on average. We therefore asked whether this distributed information could support a more robust population-level representation of trial direction.

We reasoned that if task-relevant information is reliably maintained in the dmPFC, population activity should reliably distinguish left-from right-cued trials across repeated trials. To test this, we trained a linear discriminant analysis (LDA) classifier to decode trial direction using a cross-validation (80% training, 20% testing) and compared decoding performance between correct and incorrect trials (Fig. 4F). ROC analysis showed markedly better directional discrimination on correct trials than on incorrect trials, indicating that successful task performance was associated with a stronger population-level representation of trial direction. Together, these findings indicate that, although individual dmPFC neurons exhibit only modest directional selectivity on average, directional information is robustly represented in distributed population activity.

#### Muscarinic signaling in sequential dynamics and population-level directional encoding in the dmPFC

Having established that directional information is robustly represented in distributed dmPFC population activity under baseline conditions, we next asked whether muscarinic signaling is required to maintain this representation over time. Based on our *ex vivo* findings that muscarinic receptor activation preserves effective spike transmission during repetitive thalamofrontal stimulation, we hypothesized that muscarinic receptor blockade would destabilize delay-period population dynamics and impair directional encoding in the dmPFC. We first examined the temporal stability of population activity by calculating Pearson correlation coefficients between population activity vectors across time bins and constructing temporal correlation matrices under pre-injection, saline, and atropine condition (Fig. 5). In correct trials under control conditions, population activity remained correlated across extended portions of the delay period, producing a broad band of elevated correlation around the diagonal (Fig. 5A). This temporal structure was reduced in error trials (Fig. 5B), indicating an association between successful performance and greater stability of delay-period population activity. Atropine similarly reduced the temporal stability of delay-period temporal structure even in correct trials compared with pre-injection and saline conditions (Fig. 5C-D). To quantify this effect, we measured the decay of population-vector correlation as a function of temporal separation (Δt; Fig. 5E-F). Correlation declined significantly more rapidly following atropine administration than under pre-injection or saline conditions, indicating that muscarinic receptor blockade accelerated the divergence of dmPFC population activity over time.

**Figure 5.**
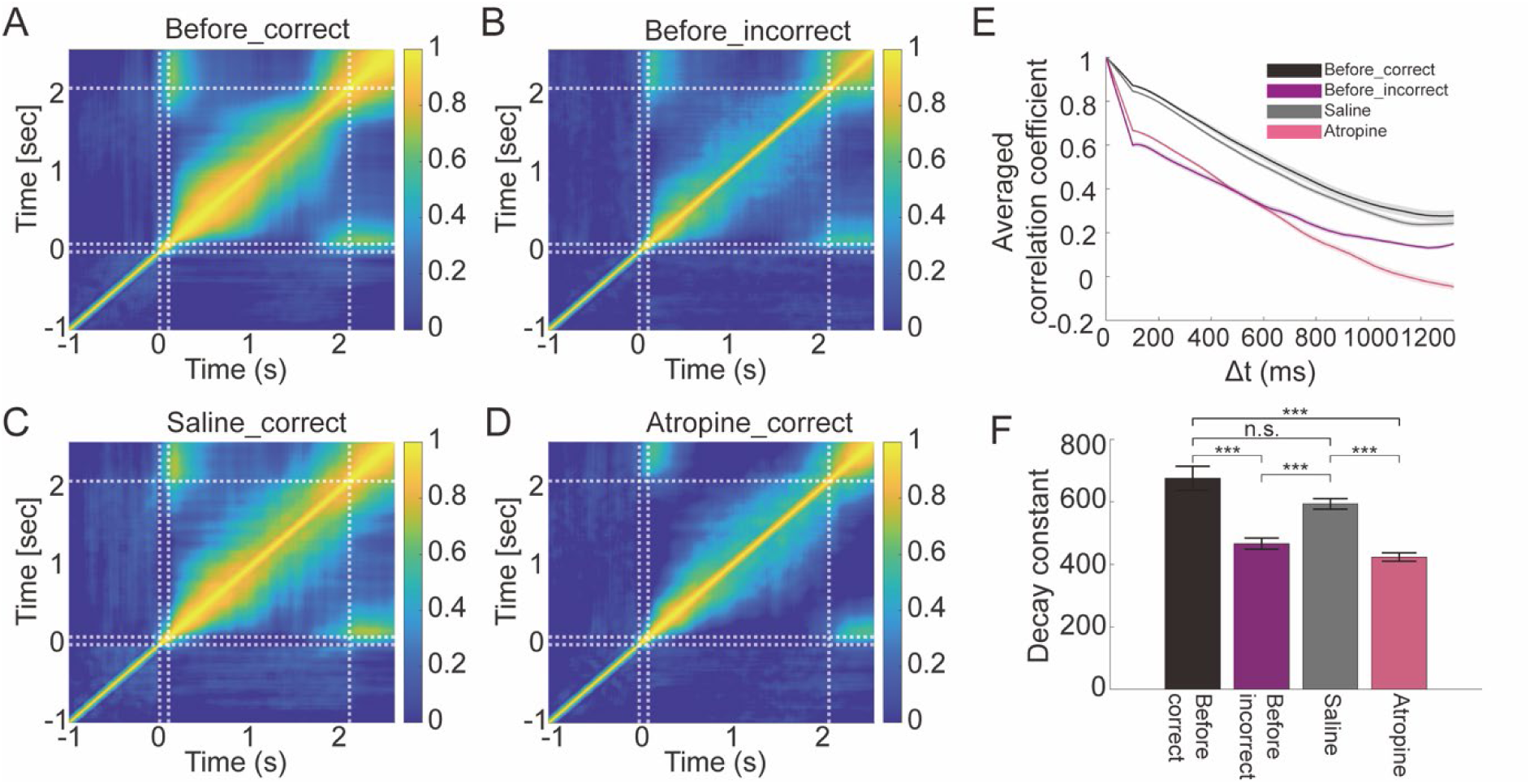
Cross-temporal stability of dmPFC population activity during STM. **(A-D)** Cross-temporal population correlation matrices showing Pearson correlation coefficients between population activity vectors across time bins aligned to auditory cue onset (t = 0) for **(A)** Correct pre-injection trials, **(B)** incorrect pre-injection trials, **(C)** correct trials following saline administration, and **(D)** correct trials following atropine administration. Vertical and horizontal dashed lines indicate cue onset and the end of the 2-s delay period. **(E)** Mean population correlation as a function of temporal separation (Δt). Shaded areas indicate SEM. **(F)** Decay constants obtained from fits to the correlation-versus-Δt curves for each condition. Smaller decay constants indicate more rapid divergence of population activity patterns over time.

We next asked whether the reduced temporal stability of population activity was accompanied by impaired population-level directional encoding. Because neurons were recorded across different sessions, we constructed pseudo-populations to estimate ensemble activity. For each iteration, 100 neurons were randomly subsampled to generate population vectors, and trial direction was decoded using a cross-validated LDA classifier (80% training, 20% testing; Methods). Decoder performance, quantified by auROC, was significantly reduced following atropine administration compared with control conditions (Fig. 6A). Thus, muscarinic receptor blockade impaired the population-level representation of trial direction in the dmPFC, accompanying the reduced temporal stability of delay-period activity. Because saline administration did not significantly alter decoding performance relative to the pre-injection condition, subsequent analyses focused on the pre-injection and atropine conditions (Fig. 6B).

**Figure 6.**
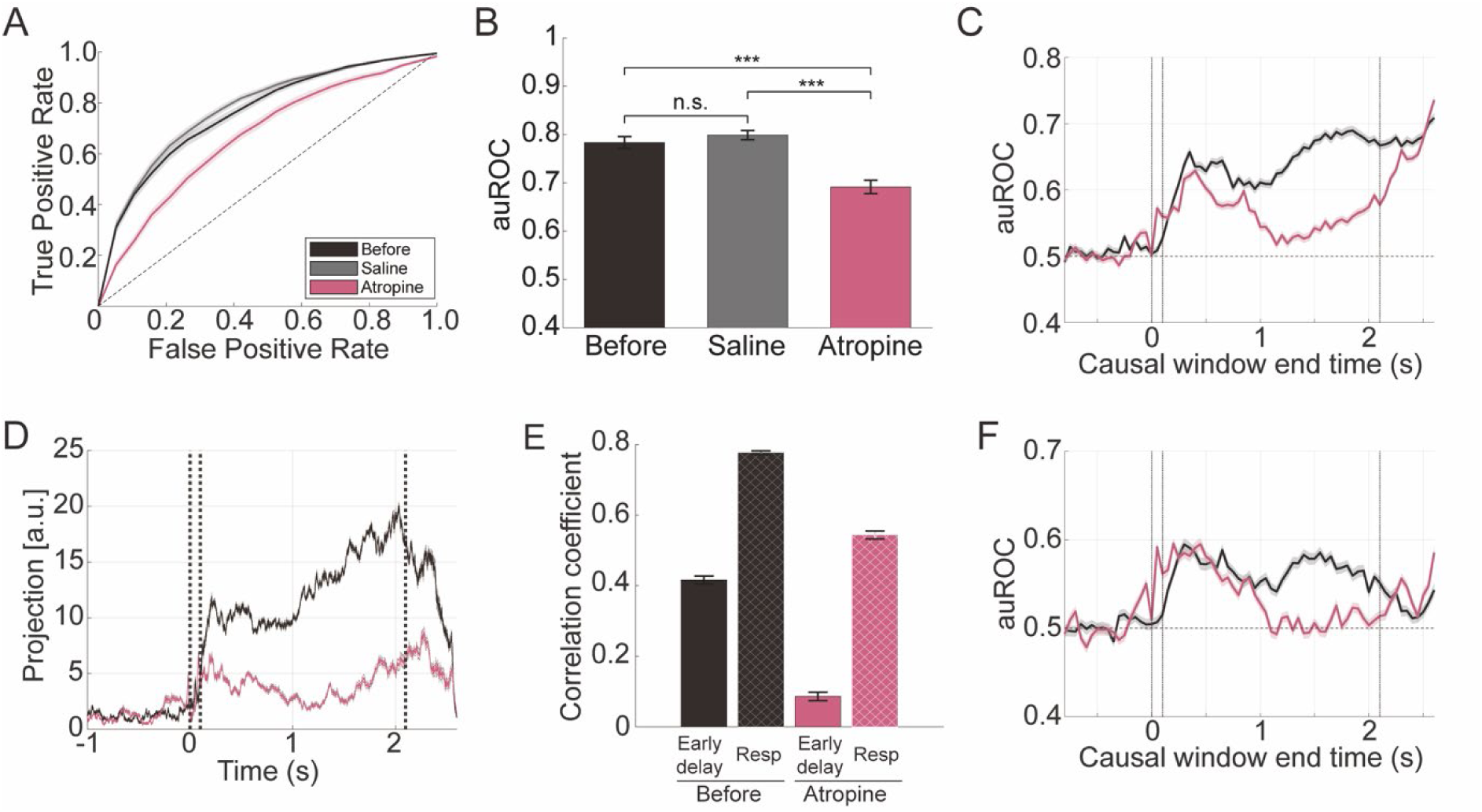
Muscarinic regulation of population-level directional representations in the dmPFC. **(A)** Receiver operating characteristic (ROC) curves for population-level decoding of trial direction using a cross-validated linear discriminant analysis (LDA) applied to pseudo-populations of 100 neurons under pre-injection (black), saline (gray), and atropine (pink) conditions. **(B)** Decoder performance quantified by auROC across pre-injection, saline, and atropine conditions, \*\*\**p*<0.001. **(C)** Time-resolved directional decoding under pre-injection (black) and atropine (pink) conditions. Decoder performance was calculated using 200-ms causal windows aligned to cue onset (t = 0). Vertical lines indicate cue onset/offset and the end of the delay period; the horizontal dashed line indicates chance performance (auROC = 0.5). **(D)** Directional separation along the delay coding direction (CD) under pre-injection (black) and atropine (pink) conditions. The CD axis was defined from delay activity in training trials, and held-out population activity was projected onto this fixed axis throughout the trial. Projection values were normalized relative to trial-shuffled distributions. **(E)** Similarity of late-delay population activity (0.5 s preceding response onset) to early-delay activity (solid bars) and response-period activity (patterned bars) under pre-injection (black) and atropine (pink) conditions, quantified by Pearson correlation coefficients. **(F)** Time-resolved directional decoding after excluding response-modulated neurons under pre-injection (black) and atropine (pink) conditions.

We next asked whether the atropine-induced impairment in population-level directional representation emerges progressively during the delay period. Given that systemic atropine did not impair performance in zero-delay trials (Fig. 3H, I), we hypothesized that mAChR blockade would preferentially affect the maintenance of directional information over time rather than its initial establishment. We therefore examined the temporal evolution of directional decoding using the same pseudo-population approach, calculating decoder performance in 200-ms time bins with a cross-validated LDA classifier (Methods). Consistent with this prediction, atropine produced a comparable initial increase in decoding performance following cue onset, but decoding accuracy subsequently declined and failed to reach the sustained levels observed under control conditions (Fig. 6C). Thus, muscarinic receptor blockade relatively spared the initial directional representation but progressively impaired its maintenance across the delay period.

We next asked whether the directional population representation established during the early delay was maintained as the delay progressed. Using a cross-validated coding-direction (CD) analysis, we defined a population axis capturing the difference between left- and right-trial activity during early-delay in the training trials. Held-out population activity was then projected onto this fixed axis throughout the trial (Methods). CD values were normalized against null distributions generated from trial-shuffled data. Unlike time-resolved decoding, which can exploit different population patterns at different time points, this analysis tests whether the coding dimension established during the early delay persists over time. In the pre-injection condition, separation of left- and right-trial activity along the early-delay coding direction remained elevated throughout the delay (Fig 6D, black). Under atropine, directional separation was also evident during the early delay but subsequently declined (Fig 6D, pink), indicating that the population representation established early in the delay was not maintained along the same coding dimension.

The delayed emergence of these neural deficits predicted that behavioral performance should remain relatively preserved when the maintenance interval was short. We therefore examined task performance across short delays of up to 0.8 s following systemic atropine administration. Atropine did not significantly impair performance at these short delays (Extended Data Fig. 4G), in contrast to the pronounced impairment observed at longer delays. Thus, the temporal emergence of the behavioral deficit paralleled the progressive disruption of task-relevant population representations in the dmPFC.

Despite the progressive decline in directional representation during the delay, both LDA decoding performance and directional separation along the CD increased again toward the onset of the response period. Because these analyses were performed on correct trials, trial direction was necessarily coupled to the upcoming directional licking response, raising the possibility that the late recovery reflected emerging response-related activity rather than restoration of the earlier delay-period representation. To examine this possibility, we compared late-delay population activity (the 0.5 s preceding response onset) with activity during the first 0.5 s of the delay period and the first 0.5 s of the response period. Late-delay population activity was significantly more strongly correlated with response-period activity than with early-delay activity (Fig. 6E), indicating that the population state had begun to transition toward a response-related configuration before response onset. We next identified response-modulated neurons (Methods) and repeated the time-resolved LDA analysis after excluding these neurons and sampling 100 neurons from the remaining pool. Excluding response-selective neurons substantially attenuated the late recovery in directional decoding (Fig. 6F). Together, these results indicate that the increase in directional discriminability toward the end of the delay was largely associated with the emergence of response-related population activity rather than restoration of the earlier delay-period representation.

Given that repetitive thalamofrontal input became progressively less effective at driving dmPFC neurons because of short-term depression, whereas mAChR activation maintained effective spike transmission during repetitive stimulation (Fig. 3), we asked whether muscarinic blockade increasingly disrupted direction-dependent neuronal activity as the delay progressed. Atropine reduced the fraction of directionally selective neurons during both periods, with a significantly greater relative reduction during the late delay (Fig. 7A-B). We next quantified the magnitude of directional firing-rate modulation as the absolute difference in normalized firing rate between right- and left-choice trials (|RR−LL|). Atropine reduced directional firing-rate modulation during the early delay, with the largest reduction observed during the late delay (Fig. 7C-D).

**Figure 7.**
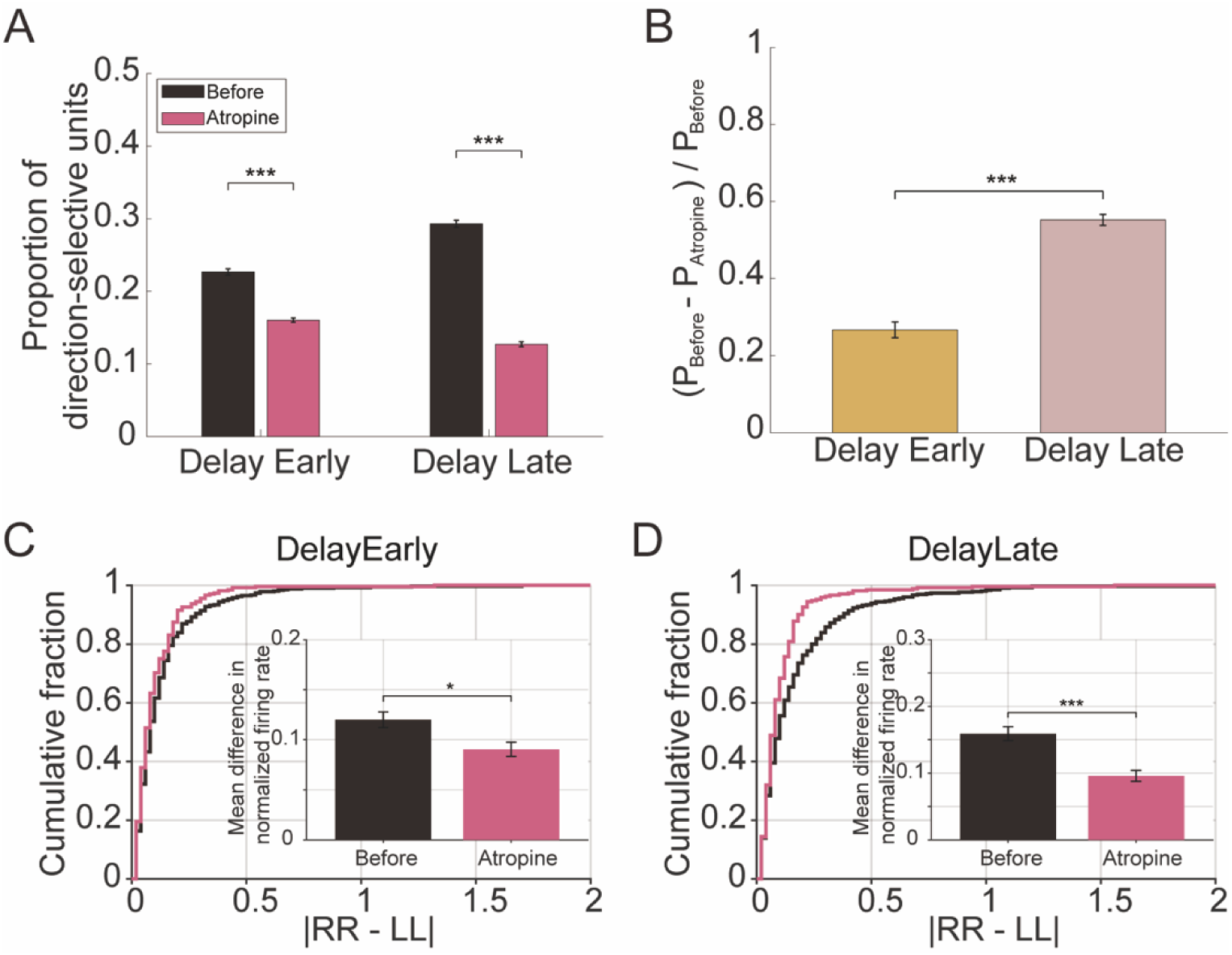
Muscarinic modulation of single-neuron directional selectivity in the dmPFC during the delay period. **(A)** Proportion of direction-selective units during the early and late delay periods under pre-injection (Before, black) and atropine (pink) conditions (\*\*\**p*<0.001). **(B)** Relative reduction in the proportion of direction-selective units following atropine treatment [(P_Before_ – P_Atropine_)/P_Before_] compared between early and late delay periods (\*\*\**p*<0.001). **(C)** Cumulative distribution of directional firing-rate modulation during the early delay, quantified as the absolute difference in normalized firing rate between right- and left-choice trials (|RR-LL|), under pre-injection (black) and atropine (pink) conditions. Inset, mean |RR-LL| under each condition (\**p*<0.05). **(D)** Same as (C), but for the late delay period (***p<0.001).

Together, these findings indicate that muscarinic signaling supports the maintenance of task-relevant dmPFC activity as the delay progresses. Although directional representations were established during the early delay under muscarinic receptor blockade, they became progressively less stable and less discriminable over time, accompanied by reductions in both the proportion of directionally selective neurons and the magnitude of direction-dependent firing-rate modulation. This temporal profile is consistent with the progressive weakening of repetitive thalamofrontal excitation caused by short-term synaptic depression and with the ability of muscarinic signaling to preserve postsynaptic responsiveness despite that weakening, as observed *ex vivo*.

## DISCUSSION

In this study, we investigated how thalamofrontal communication is sustained during STM despite the strong short-term depression of thalamofrontal synapses. We found that muscarinic cholinergic signaling supports the maintenance of directional information and task-relevant population dynamics in the dmPFC, with its contribution becoming particularly evident as the memory delay progresses. Together, our findings identify cholinergic regulation of thalamofrontal transmission as a mechanism linking the synaptic properties of this pathway to the maintenance of cortical representations during STM.

MD provides a major source of excitatory drive to the prefrontal cortex and is critical for sustaining cortical dynamics during cognitive tasks ^18,9^. During STM, even temporally restricted inhibition of MD→dmPFC projections impairs memory performance ^10^. Our results, however, reveal a substantial biophysical constraint on this pathway: MD→dmPFC synapses exhibit profound short-term depression (Fig. 1). Consequently, repetitive thalamic activity produces progressively weaker postsynaptic responses, effectively imposing a low-pass filtering property on thalamofrontal transmission ^45,46^. Thus, without cholinergic enhancement of postsynaptic excitability, short-term depression would be expected to progressively constrain the ability of sustained MD activity to drive dmPFC neurons.

Our *ex vivo* experiments demonstrate that muscarinic receptor activation increases the intrinsic excitability of dmPFC pyramidal neurons (Fig. 3), consistent with the suppression of potassium conductance and consequent increases in membrane resistance described previously as selective ‘attention-tuning’ ^57^. Increased membrane resistance enhances the voltage response produced by a given synaptic current, allowing dmPFC neurons to reach spike threshold even as thalamofrontal synaptic currents progressively weaken during repetitive activity. In this framework, ACh acts as a permissive gate for thalamofrontal transmission: rather than preventing short-term synaptic depression, muscarinic signaling adjusts postsynaptic excitability to maintain effective excitation transfer despite it. Such state-dependent regulation of input–output gain provides a potential cellular mechanism for the proposed role of cholinergic signaling in attentional tuning ^59^, by determining whether otherwise subthreshold afferent activity can effectively recruit cortical neurons.

A key advantage of cholinergic regulation of thalamofrontal transmission is its temporal and behavioral specificity ^60,61,31,62^. Cortical ACh rose rapidly during task engagement, remained elevated throughout the memory delay, and returned toward baseline within seconds after task completion, while sustained elevation was absent in naïve animals (Fig. 2). Such dynamics are well suited for transiently enhancing thalamofrontal transmission when task-relevant activity must be maintained. Consistent with this idea, muscarinic receptor activation increased dmPFC neuronal excitability and enhanced responsiveness to ongoing thalamic input. ACh may therefore act as a temporally restricted gain signal that promotes the persistence of task-relevant prefrontal representations without itself providing the information being represented.

Other neuromodulatory systems may complement cholinergic regulation by acting at distinct sites and on different components of synaptic transmission. For example, noradrenaline suppresses thalamocortical transmission through presynaptic α2A-adrenoceptors, reducing transmitter release probability and thereby altering short-term synaptic dynamics ^63^. Dopamine D1/D5 receptor activation in the PFC, in contrast, enhances the postsynaptic NMDA component of excitatory transmission and promotes temporal summation during repetitive synaptic activity ^64^. Thus, neuromodulators may support effective information transfer during repetitive activity through complementary regulation of presynaptic release dynamics, postsynaptic integration, and intrinsic excitability.

Increasing postsynaptic responsiveness also raises the question of how task-relevant signals can be enhanced without indiscriminately amplifying background activity. ACh differentially regulates cortical interneuron classes ^65,66^, and cholinergic modulation can improve neuronal discriminability and reduce interneuronal correlations in visual cortex ^67,68^. Local inhibitory circuitry may therefore complement cholinergic enhancement of pyramidal-cell excitability to preserve coding fidelity, although whether this occurs in dmPFC during STM remains unknown.

An important implication of ACh-mediated enhancement of postsynaptic excitability is that thalamofrontal communication should become increasingly dependent on cholinergic modulation as synaptic depression accumulates during the delay. Early in the delay, before the effects of repetitive synaptic depression have accumulated, directional representations may remain relatively resistant to muscarinic blockade; as the delay proceeds, however, progressively weakened thalamofrontal input should place increasing demands on cholinergic enhancement of postsynaptic excitability (Fig. 6). This effect can be understood in terms of the nonlinear input–output relationship of cortical neurons: as synaptic drive declines, dmPFC neurons may move toward a near-threshold, lower-gain regime in which differences between competing inputs are less effectively translated into differences in spiking. By increasing neuronal excitability, cholinergic signaling may help maintain prefrontal neurons within an operating range that preserves the effective representation of task-relevant information. Consistent with this view, nucleus basalis stimulation in primates increased prefrontal delay-period activity and improved working-memory performance, even though individual neurons became less spatially selective ^69^. Modeling suggested that increased excitability could improve memory by reducing drift in the population representation, despite this broader tuning ^70^. Thus, cholinergic modulation need not simply sharpen neuronal tuning, but may instead stabilize population representations by regulating the operating state of the cortical network. In our experiments, atropine largely spared directional decoding during cue presentation and the early delay but progressively impaired population coding and direction-dependent neuronal activity as the maintenance interval increased (Fig. 6). Behavioral performance showed a similar temporal dependence: short-delay performance (∼800 ms) was relatively preserved, whereas longer delays produced marked impairment (Extended Data Fig. 4). Together, these findings suggest that muscarinic signaling is more important for sustaining task-relevant information over time than for establishing the initial representation.

Beyond its cellular mechanism, the temporal specificity of this cholinergic effect also suggests a distinct role within broader frameworks of memory gating. Models of working memory have emphasized input gates that regulate which information enters working memory and output gates that determine which maintained representations influence subsequent processing or behavior ^71^. This distinction provides a useful framework for interpreting our STM task. Muscarinic blockade largely spared the initial establishment of directional representations but impaired their persistence as the delay progressed, suggesting that cholinergic modulation of thalamofrontal transmission may provide a maintenance-oriented form of gating that regulates whether an established representation continues to receive sufficient excitatory support over time.

Several limitations should be considered. First, we could not manipulate muscarinic signaling with sufficient temporal precision to determine whether the initial cue-evoked cholinergic response and the sustained delay-period elevation make distinct contributions to STM. The ACh signal evoked by the cue persisted on a timescale comparable to the delay itself (Fig. 2), and suppression of subsequent cholinergic release would not necessarily terminate receptor activation or downstream muscarinic signaling already initiated by the initial ACh elevation. Thus, temporally restricted inhibition of cholinergic terminals alone may be insufficient to isolate the contribution of individual phases of the cholinergic response. Approaches allowing more rapid and reversible control of muscarinic receptor signaling, potentially combined with direct monitoring of extracellular ACh, will be required to resolve the respective contributions of phasic and sustained cholinergic activity. Second, atropine was administered systemically, and the behavioral and population-level effects therefore cannot be attributed exclusively to muscarinic signaling within the dmPFC. The preservation of zero-delay performance argues against a gross sensory or motor deficit, but muscarinic blockade in other cortical or subcortical regions may nevertheless contribute to the STM impairment. Local or cell-type-specific manipulations of muscarinic signaling in the dmPFC will be required to establish its regional and cellular specificity. Third, although our *ex vivo* experiments identify increased postsynaptic excitability as an important mechanism by which muscarinic signaling preserves effective responses to repetitive thalamofrontal input, the extent to which this mechanism specifically supports thalamofrontal transmission *in vivo* remains unresolved. Increased dmPFC excitability could also enhance recurrent corticocortical interactions or other afferent inputs that contribute to memory maintenance. Thus, the progressive loss of directional information under muscarinic blockade cannot be attributed exclusively to impaired thalamofrontal transmission. Pathway-specific manipulations of MD→dmPFC input, together with measurements of thalamofrontal synaptic efficacy during behavior, will be important for determining how strongly the cellular mechanism identified ex vivo contributes to maintenance of dmPFC representations in vivo.

In conclusion, our study identifies a cholinergic gating mechanism that helps overcome a biophysical constraint imposed by short-term depression at thalamofrontal synapses. By increasing postsynaptic excitability through muscarinic receptor activation, cholinergic signaling enables dmPFC neurons to remain responsive to progressively weakened thalamic input, thereby supporting the maintenance of task-relevant population representations during STM. This mechanism links the synaptic dynamics of thalamofrontal transmission to the maintenance of task-relevant prefrontal representations and provides a framework for understanding how alterations in cholinergic signaling reported in aging, Alzheimer’s disease, and schizophrenia may contribute to memory dysfunction ^72–75^.

## Supporting information

https://doi.org/10.5281/zenodo.23724877

## Acknowledgments

We thank members of the Rah laboratory for helpful discussions. The authors acknowledge the use of ChatGPT (OpenAI) for assistance in correcting grammatical errors and improving text clarity. All content was subsequently reviewed and edited by the authors, who take full responsibility for the final manuscript.

## Data, code, and materials availability

Source data, associated datasets, and custom MATLAB code are publicly available through zenodo: https://doi.org/10.5281/zenodo.23724877 and github: https://github.com/JinmoKim08-bcc/KimKimEtAl_muscarinic_neuropixels.

