## Supplementary material for "Muscarinic signaling sustains prefrontal representations during short-term memory": https://doi.org/10.5281/zenodo.23724877

Table 1

Action potential-laser duration reliability of MD-h134R neurons

| | **Duration** | **0.5 ms** | **1 ms** | **3 ms** | **5 ms** |
|---|---|---|---|---|---|
| **5 Hz** | **Mean** | 40.00 | 73.33 | 100.00 | 88.00 |
| | **Std. Deviation** | 42.43 | 35.02 | 0.00 | 26.83 |
| | **Std. Error of Mean** | 18.97 | 14.30 | 0.00 | 12.00 |
| **10 Hz** | **Mean** | 40.00 | 76.67 | 100.00 | 100.00 |
| | **Std. Deviation** | 42.43 | 40.82 | 0.00 | 0.00 |
| | **Std. Error of Mean** | 18.97 | 16.67 | 0.00 | 0.00 |
| **20 Hz** | **Mean** | 36.00 | 86.67 | 88.00 | 92.00 |
| | **Std. Deviation** | 35.78 | 20.66 | 26.83 | 18.89 |
| | **Std. Error of Mean** | 16.00 | 8.43 | 12.00 | 8.00 |

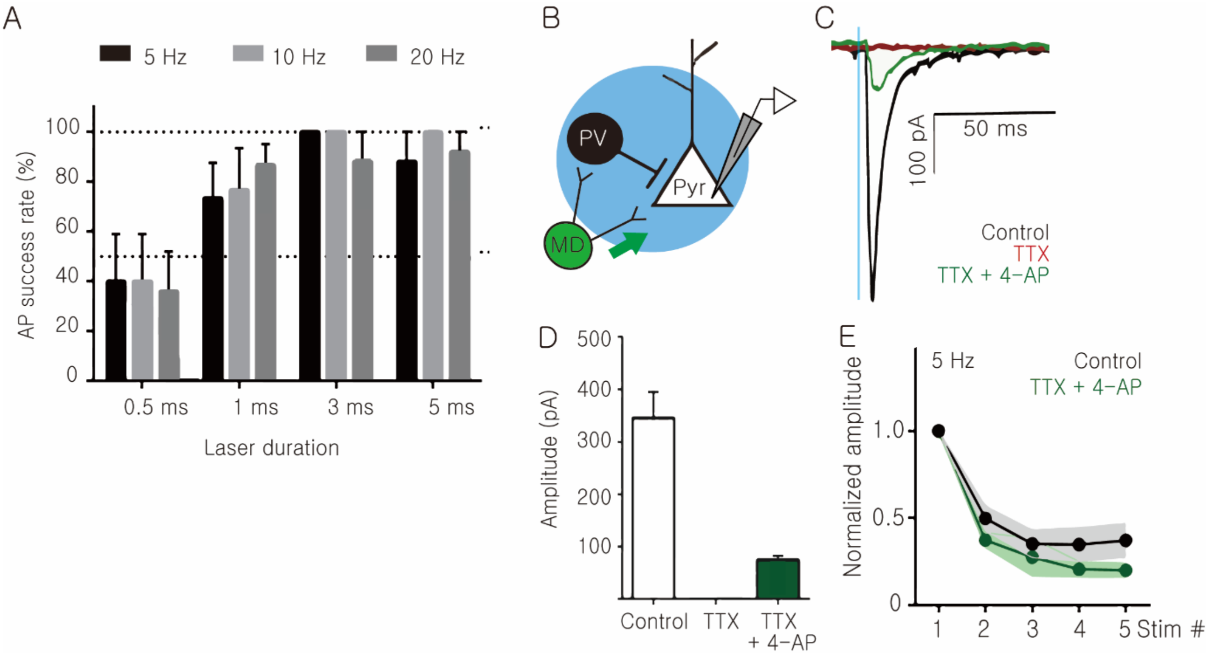


**Supplementary figure 1 H134R supports action potential generation at stimulation frequencies of up to 20 Hz and exhibits short-term dynamics of monosynaptic EPSCs**

(A) H134R-expressing MD neurons are able to generate action potentials by high-frequency stimulus (n=5)

(B) Schematic image of the recording monosynaptic thalamocortical EPSC with 0.5 μM Tetrodotoxin (TTX) and 100 μM 4-Aminopyriding (4-AP).

(C) Sample traces of monosynaptic EPSC. MD-evoked EPSCs (black) were totally blocked by 0.5 μM TTX application (red) and recovered by following 100 μM 4-AP treatment (green).

(D) Comparison between MD-evoked EPSC amplitude on dACC pyramidal neurons, with TTX, or with TTX + 4-AP subsequently.

(E) Monosynaptic EPSC depresses rapidly in similar to the original EPSC at 5 Hz, 10 Hz, and 20 Hz ($p>0.05$, paired t-test).

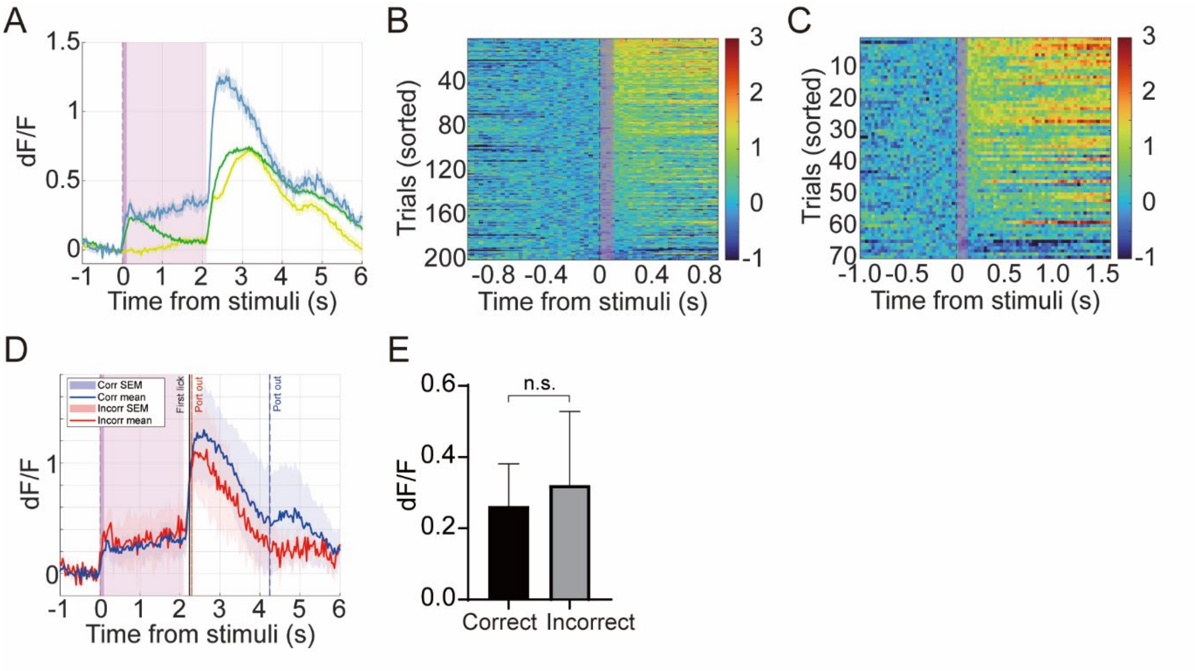


**Supplementary figure 2 ACh dynamics during the delay period and their relationship to behavioral performance**

(A) Average dF/F timecours aligned to stimulus onset. The shaded area indicates the delay period. Traces show trained mice (blue), naïve mice with an auditory cue (green), and naïve mice without an auditory cue (yellow).

(B-C) Heatmaps showing single-trial dF/F dynamics from representative session aligned to stimulus onset for delay lenghs of (B) 0.8 s and (C) 1.5 s. Trials are sorted by the onset latency of the dF/F signal increase.

(D) Average dF/F dynamics for correct and incorrect trials across sessions.

(E) Comparison of mean dF/F during the delay period between correct and incorrect trials, showing no significant difference.

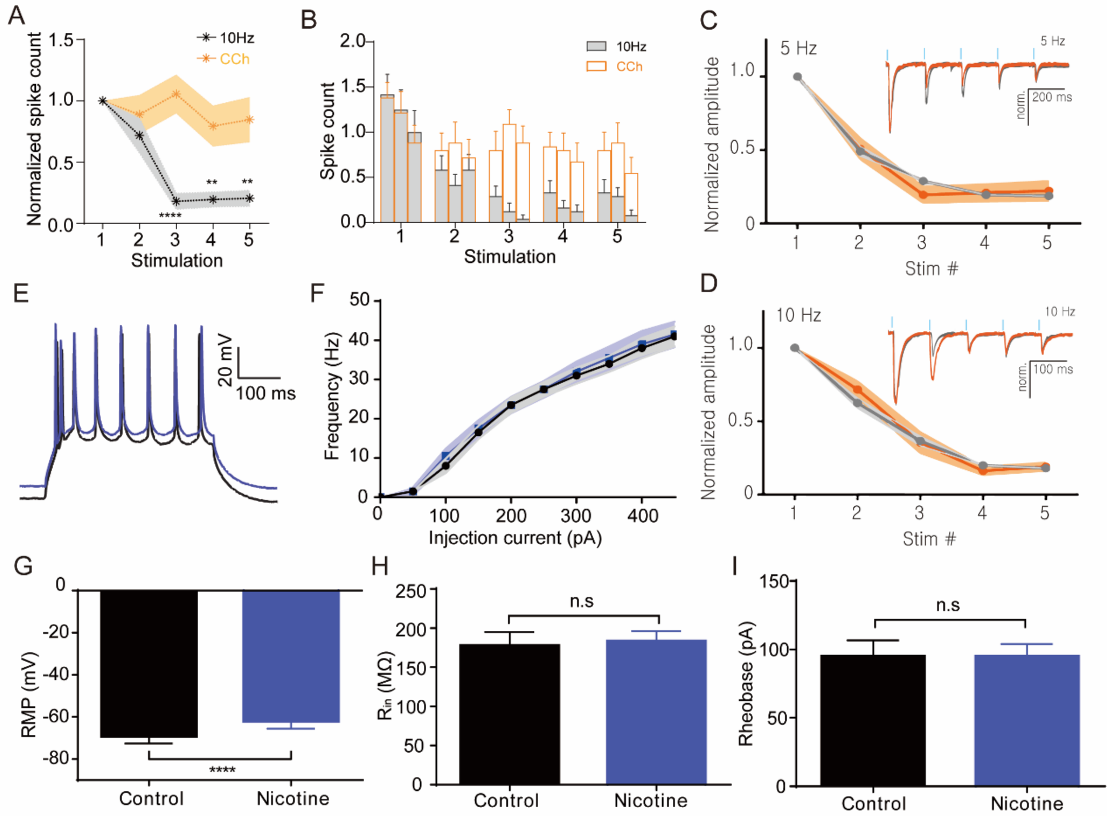


**Supplementary figure 3 Cholinergic activation enhances thalamofrontal transmission by increasing spike probability and intrinsic excitability through mAChR**

(A) Normalized spike count during high-frequency stimulation of MD axons in dmPFC pyramidal neurons under control conditions (black) and in the presence of CCh. CCh significantly increased the probability of spike generation.

(B) Summary of spike counts across stimulation trials from (A), showing a higher spike probability with CCh

(C-D) Normalized amplitudes of excitatory postsynaptic responses (EPSPs) at TF synapses during (C) 5 Hz and (D) 10 Hz stimulation train, in control and CCh conditions. Inset: Representative traces. Laser stimulation is shown as a blue vertical line.

(E) Representative membrane potential traces in response to current injection under control and nicotine (100 μM) conditions. Nicotine induced a small but significant depolarization.

(F) F-I relationship showing firing frequency as a function of injected current under control and nicotine conditions

(G-I) Effects of nicotine on intrinsic excitability parameters, including (G) resting membrane potential, (H) input resistance, and (I) rheobase current.

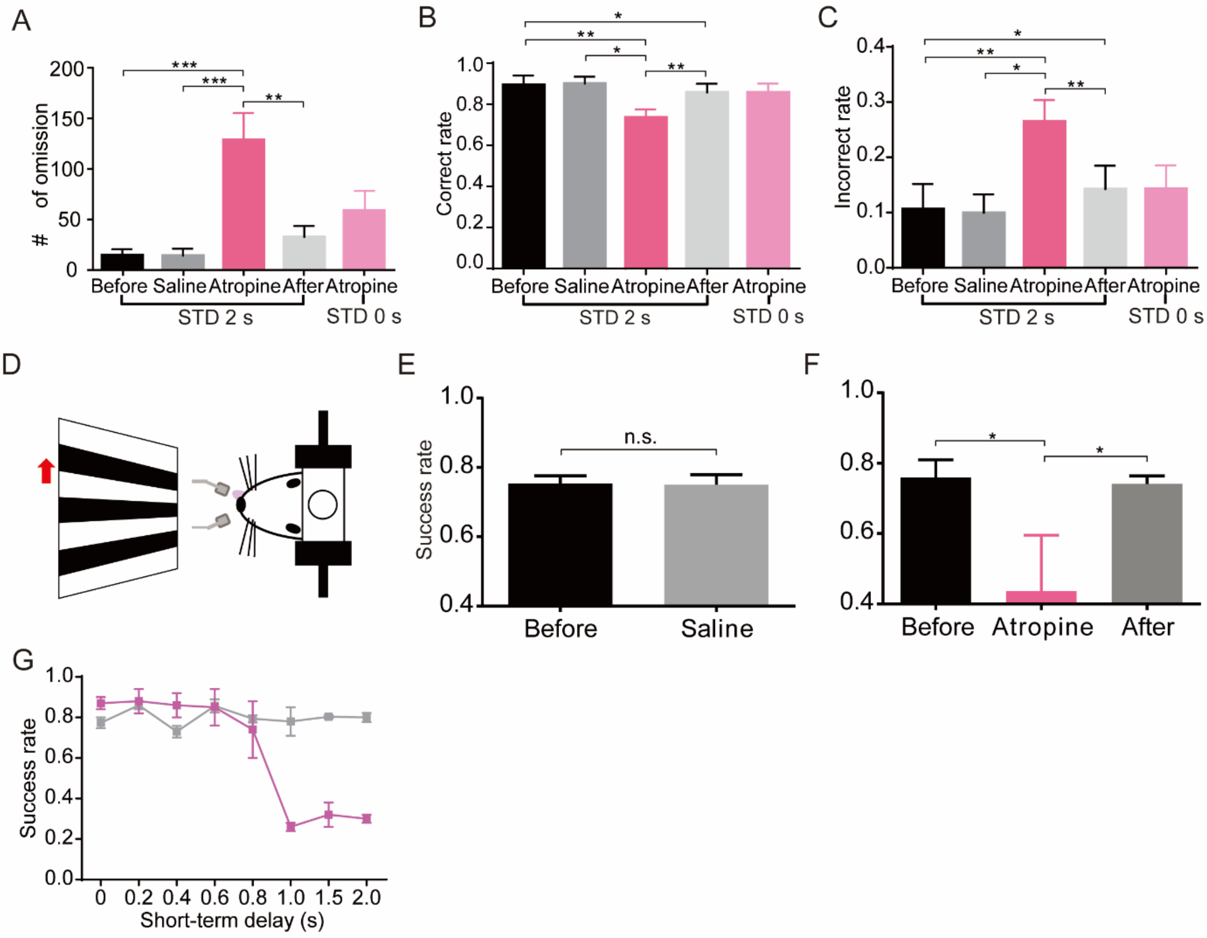


**Supplementary figure 4 Atropine impairs short-term memory performance by disrupting muscarinic signaling in the dmPFC, independent of sensory modality and task engagement**

(A) Increased number of omission trials by atropine administration.

(B) Proportion of correct trials before and after atropine administration. Atropine significantly reduced response accuracy during the short-term memory tasks (STD 2 s) but not during the discrimination tasks (STD 0 s)

(C) Proportion of incorrect trials. Atropine significantly increased the proportion of incorrect trials during STM tasks, but not during the discrimination tasks.

(D) Schematic of the visual STM task with bilateral dmPFC microinfusion. Mice were trained to memorized the direction of drifting grating motion (red arrow) and report the direction with a licking response.

(E-F) performance in the visual STM task following bilateral dmPFC infusion of saline or atropine. Atropine significantly reduced the success rate compared to saline.

(G) Time course of success rate during the short-term delay period following saline (gray) or atropine (pink) injection (i.p.). Performance remained stable following saline infusion, whereas atropine-treated mice maintained performance up to a delay of 0.8 s, followed by a marked decrease at delay of 1.0 s and longer.

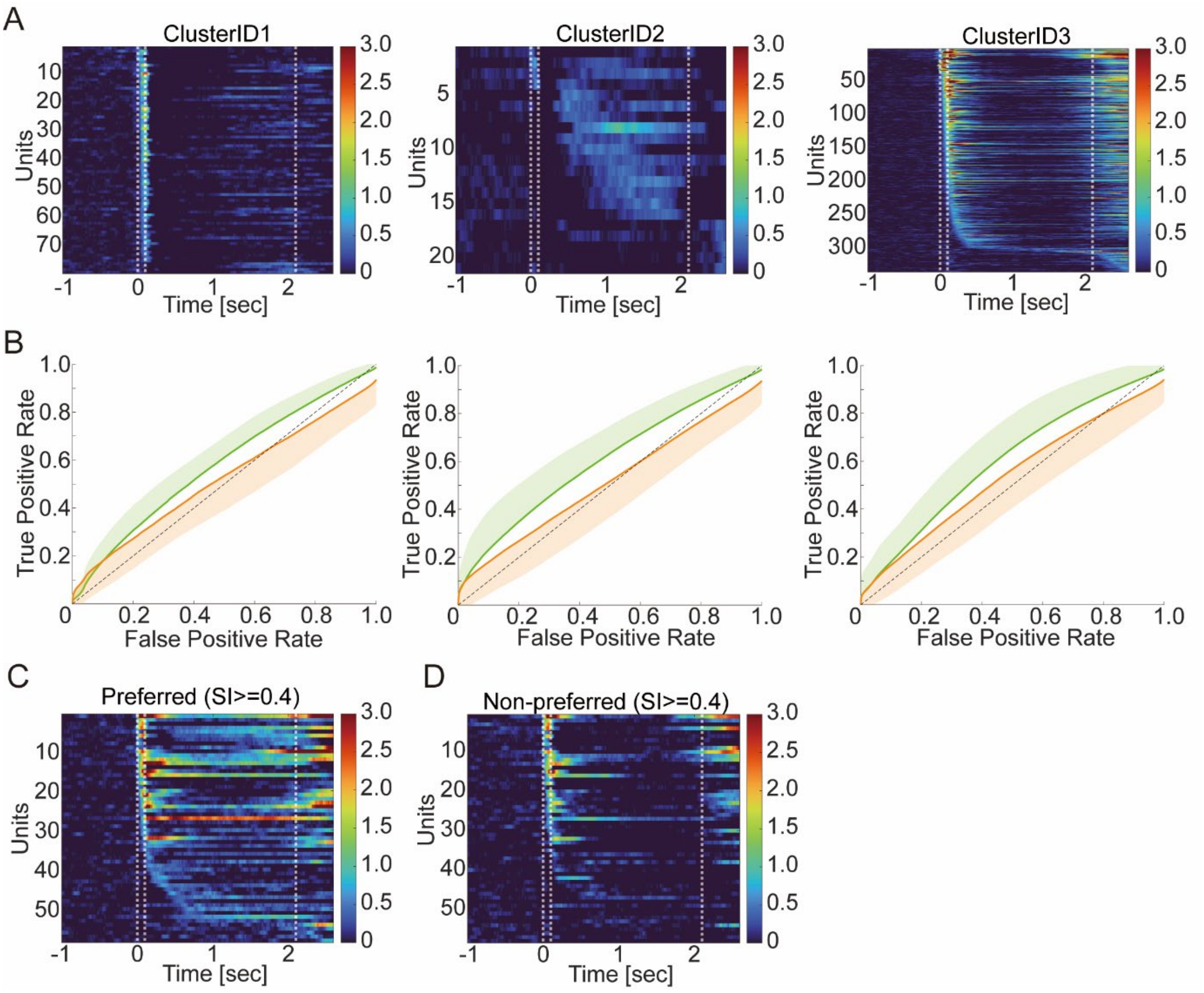


**Supplementary figure 5 Unsupervised clustering reveals three major response classes in dmPFC neurons, with comparable directional selectivity across classes but a subset of strongly selective neurons**

(A) Representative peri-event time raster plots of three neuronal response classes identified by unsupervised clustering. ClusterID1 shows cue-responsive neurons (left), ClusterID2 shows neurons with sustained delay activity (middle), and ClusterID3 shows neurons with ramping activity preceding the response (right).

(B) ROC curves for directional selectivity within each response class (ClusterID1-3). True positive rate is plotted against false positive rate.

(C-D) Peri-event time heatmaps of directionally selective neurons (SI ≥ 0.4) in the preferred (C) and non-preferred (D) direction.
